# The MOM1 complex recruits the RdDM machinery via MORC6 to establish *de novo* DNA methylation

**DOI:** 10.1101/2023.01.10.523455

**Authors:** Zheng Li, Ming Wang, Zhenhui Zhong, Javier Gallego-Bartolomé, Suhua Feng, Yasaman Jami-Alahmadi, Xinyi Wang, James Wohlschlegel, Sylvain Bischof, Jeffrey A. Long, Steven E. Jacobsen

**Author notes:** These authors contributed equally (Zheng Li, Ming Wang, Zhenhui Zhong).

## Abstract

MOM1 is an *Arabidopsis* factor previously shown to mediate transcriptional silencing independent of major DNA methylation changes. Here we found that MOM1 localizes with sites of RNA-directed DNA methylation (RdDM). Tethering MOM1 with artificial zinc finger to unmethylated *FWA* promoter led to establishment of DNA methylation and *FWA* silencing. This process was blocked by mutations in components of the Pol V arm of the RdDM machinery, as well as by mutation of *MORC6*. We found that at some endogenous RdDM sites, MOM1 is required to maintain DNA methylation and a closed chromatin state. In addition, efficient silencing of newly introduced *FWA* transgenes was impaired by mutation of MOM1 or mutation of genes encoding the MOM1 interacting PIAL1/2 proteins. In addition to RdDM sites, we identified a group of MOM1 peaks at active chromatin near genes that colocalized with MORC6. These findings demonstrate a multifaceted role of MOM1 in genome regulation.

## Introduction

Transcriptional silencing is critical to keep transposable elements and DNA repeats under control in eukaryotic genomes. The process of transcriptional silencing involves several elaborate mechanisms involving many proteins as well as DNA methylation and histone modifications^1,2^. In *Arabidopsis*, the *MORPHEUS’ MOLECULE1* (*MOM1*) gene, which was originally identified with the phenotype of reactivation of a DNA-methylated and silenced hygromycin-resistance transgene in the *mom1* mutant^3^, is a distinct component of the transcriptional silencing machinery. In the *mom1* mutant, a set of transposable elements, mainly located in pericentromeric regions^4–6^, is robustly activated without major alteration in DNA methylation patterns^5,7,8^. In addition, no obvious visible decompaction of heterochromatin at chromocenters was observed in the *mom1* mutant^9–11^. The mechanism of MOM1 mediated silencing remains elusive.

*MOM1* encodes a large protein (2001 amino acids) with sequence homology to the ATPase domain of SWI2/SNF2 family proteins^3^. However, this SNF2 homology sequence is largely dispensable for MOM1’s silencing function^12^. Instead, the Conserved MOM1 Motif 2 (CMM2) domain, which is conserved among MOM1 orthologs, is required for the silencing function of MOM1^12^. The CMM2 domain of MOM1 multimerizes with itself and interacts with two PIAS (PROTEIN INHIBITOR OF ACTIVATED STAT)-type SUMO E3 ligase-like proteins, PIAL1 and PIAL2^5,13^. The *pial1 pial2* double mutant phenotype highly resembles the endogenous TE de-repression phenotype of *mom1*^5^, suggesting that the PIAL proteins and the MOM1 protein function in the same pathway. However, evidence suggests that the SUMO ligase activity is not require for the transcriptional silencing by PIAL2, and the interaction of MOM1 and PIAL2 with SUMO is also not required for the silencing function of the MOM1 complex^5,14^.

RNA directed DNA Methylation (RdDM) is a plant specific pathway responsible for *de novo* DNA methylation^15^. It also assists in maintaining preexisting DNA methylation patterns together with other DNA methylation mechanisms^16^. The RdDM pathway can be divided into two arms. In the RNA POLYMERASE IV (Pol IV) arm of the RdDM pathway, SAWADEE homeodomain homolog 1 (SHH1) and CLASSY (CLSY) proteins recruit Pol IV to target sites marked by H3K9 methylation and unmethylated H3K4 to produce precursor single-stranded RNA (ssRNA) of 30-45 nucleotides in length^17–20^. RNA-directed RNA polymerase 2 (RDR2) then converts these ssRNAs into double-stranded RNAs (dsRNA), which are then processed by Dicer-like 3 (DCL3) into 24nt siRNA^21–24^. 24nt siRNA are then loaded into ARGONAUTE proteins, AGO4, AGO6 or AGO9, which then participate in the RNA POLYMERASE V (Pol V) arm of the RdDM pathway^17,25–27^. The Pol V arm of the RdDM pathway is initiated by SU(VAR)3-9 homolog 2 (SUVH2) and SUVH9 binding to methylated DNA and recruiting the DDR complex composed of the DEFECTIVE IN RNA-DIRECTED DNA METHYLATION 1 (DRD1), DEFECTIVE IN MERISTEM SILENCING3 (DMS3) and RNA-DIRECTED DNA METHYLATION1 (RDM1) proteins^28–31^. Subsequently, Pol V is recruited by the DDR complex and synthesizes non-coding RNAs which serve as scaffolds for the binding of AGO-siRNA duplexes^18,32–34^. The DNA methyltransferase enzyme DOMAINS REARRANGED METHYLTRANSFERASE 2 (DRM2) is then recruited to methylate target DNA^35^.

RNA-seq analysis shows that the majority of up-regulated genes and TEs in the *mom1* mutant and in the *nrpe1* mutant (mutant of the largest subunit of Pol V) do not overlap^5,6^. In addition, some genes are only significantly up-regulated in the *mom1 nrpe1* double mutant^6^, and a mutant allele of *nrpe1* was identified in a screen for enhancers of the de-repression of a transgenic luciferase reporter in the *mom1* background^6^. These studies suggest that, although MOM1 mediated transcriptional silencing and RdDM function as two different pathways, they also can act cooperatively to silence some endogenous and transgene targets.

The Arabidopsis Microrchidia (MORC) proteins were discovered as additional factors required for gene silencing downstream of DNA methylation^36^. In addition, MORCs associate with components of the RdDM pathway, are loaded onto sites of RdDM and are needed for the efficiency of RdDM maintenance at some sites^37–40^. The connection between the RdDM pathway and the MORC proteins has also been demonstrated through experiments targeting the *FWA* gene. In wild type plants, *FWA* expression is silenced in all tissues except the endosperm due to DNA methylation in the promoter^41^. In the *fwa-4* epi-mutant (*fwa*), the *FWA* gene promoter is unmethylated leading to constitutive expression of the *FWA* gene and late flowering phenotype^42^. Tethering MORC proteins to the unmethylated promoter of the *FWA* gene in the *fwa* mutant via protein fusion to an artificial zinc finger protein 108 (ZF) led to efficient methylation of the promoter via recruitment of the RdDM machinery^40,43^. In addition, mutations in the MORC proteins impair the efficient *de novo* methylation and silencing of *FWA* transgenes^40^.

Several previous studies have identified functional similarities between MORC proteins and the MOM1 complex. Multiple screens using silenced transgene reporters have identified mutations in both *MOM1* and *MORC6*^5,7^, suggesting that they are both required for maintaining the silenced state of these transgenes. Analysis of gene expression defects in mutants has shown that a significant proportion of derepressed TEs in the *morc6* mutant were also derepressed in *mom1*, while another group of TEs are uniquely derepressed only in the *mom1 morc6* double mutant^7^. Thus, investigating the relationship between the RdDM machinery, MORC proteins and the MOM1 complex should help to understand the convergence and divergence in their functions.

In this study, by performing chromatin immunoprecipitation sequencing (ChIP-seq) of the MOM1 protein and MOM1 complex components, we observed strong colocalization with the MORC6 protein and RdDM sites. Tethering of MOM1 complex components to the *FWA* promoter in the *fwa* mutant by ZF fusion led to the establishment of DNA methylation and silencing of the *FWA* gene. By transforming ZF fusions into mutants we discovered that the establishment of DNA methylation by ZF-MOM1 was not only blocked by the mutants of the downstream components of the Pol V arm of the RdDM pathway, but was also blocked in *morc6*. Furthermore, an interaction between PIAL2 and MORC6 was detected by a Yeast Two-Hybrid (Y2H) assay as well as co-immunoprecipitation (co-IP). In addition, efficient *de novo* methylation and silencing of an *FWA* transgene was impaired in the *mom1* and the *pial1/2* mutants. Consistent with the divergent function of the MOM1 complex and the RdDM pathway, the MOM1 complex was more enriched at TEs in pericentromeric region, while Pol V is more enriched at TEs in the chromosome arms. MOM1 also binds to a group of RdDM independent sites, at active and accessible chromatin. These results highlight new functions for MOM1 in genome regulation and help clarify the relationship between MOM1, MORCs and RdDM.

## Results

### MOM1 complex colocalizes with RdDM sites

Previously, it was shown that MOM1, PIAL1 and PIAL2 form a high molecular weight complex *in vivo*^5^. In addition, MOM1 Immunoprecipitation-Mass Spectrometry (IP-MS) pulled down other interactors such as AIPP3 and PHD1^5^. To comprehensively identify interacting components of the MOM1 complex, we repeated the IP-MS experiments of MOM1 protein with a 3X-FLAG epitope tag and observed that, consistent with previous reports, PIAL1, PIAL2, PHD1 and AIPP3 were pulled down (Fig. 1a and Supplementary Table 1). In addition, the MOM2 protein, which was predicted to be a non-functional homolog of MOM1, was identified in the MOM1 IP-MS (Fig. 1a and Supplementary Table 1). Previous IP-MS of the AIPP3 protein pulled down other protein components such as PHD2 (also called PAIPP2), PHD3 (also called AIPP2) and CPL2, in addition to PHD1^44–46^. To facilitate the dissection of the interacting components, we performed IP-MS with FLAG tagged MOM2, PIAL2, PHD1 and AIPP3. AIPP3 pulled down MOM1, MOM2, PIAL1, PIAL2, PHD1, as well as CPL2, PHD2 and PHD3 (Fig. 1a and Supplementary Table 1). However, MOM2, PIAL2 and PHD1 each pulled down each other, as well as the PIAL1 and MOM1 protein, but no peptides of CPL2, PHD2 and PHD3 (Fig. 1a and Supplementary Table 1). Thus, consistent with previous studies showing AIPP3 forms a complex with CPL2, PHD2 and PHD3^44–46^, AIPP3 appears to be a component of multiple protein complexes, one of which is the MOM1 protein complex.

**Fig. 1.**
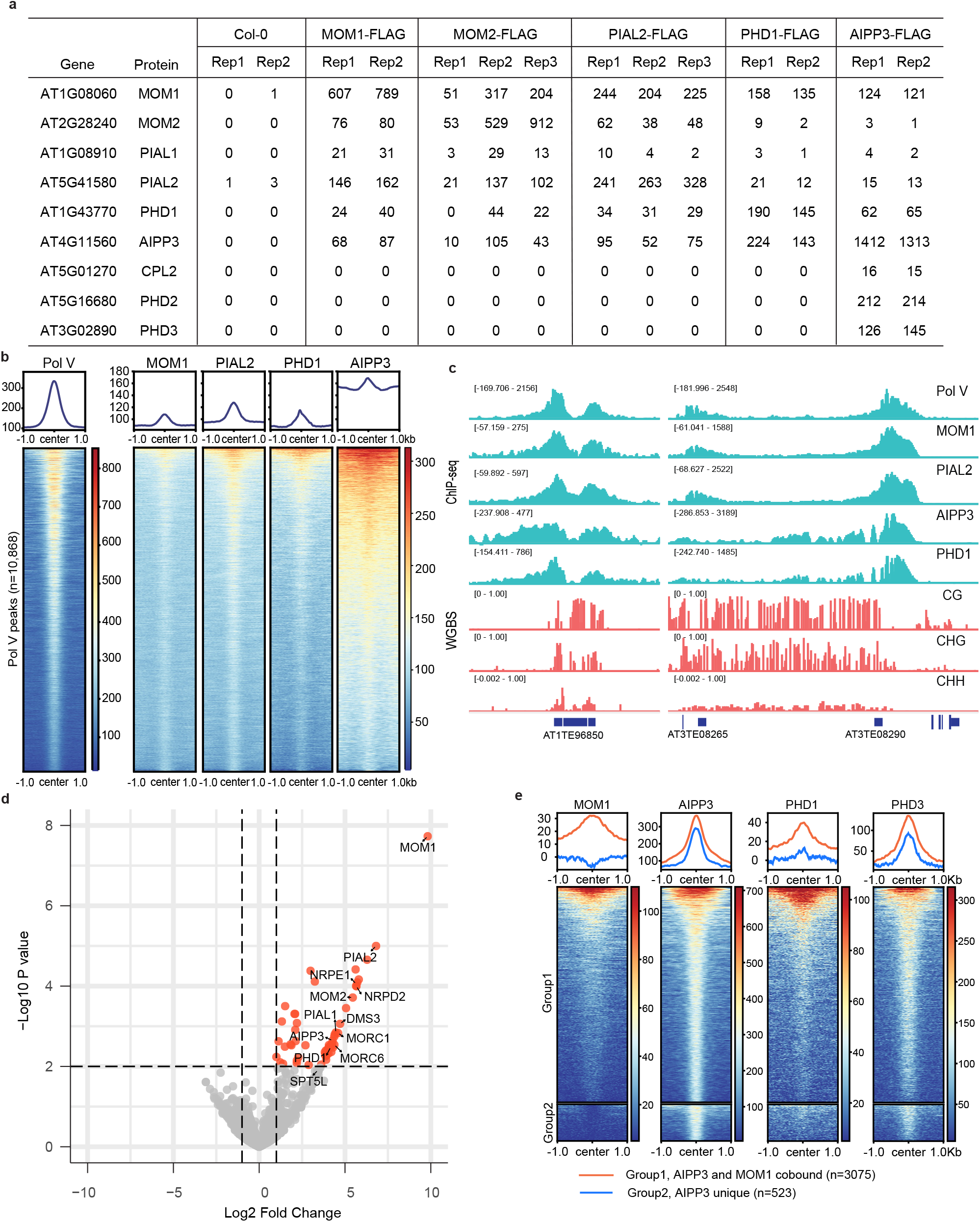
The MOM1 complex colocalizes with RdDM sites. **a**, Native IP-MS of Col-0 control and FLAG epitope tagged MOM1, MOM2, PIAL2, PHD1 and AIPP3 transgenic lines. MS/MS counts from MaxQuant output are listed. **b**, Metaplots and heatmaps representing ChIP-seq signals of Pol V, MOM1-Myc, PIAL2-Myc, PHD1-FLAG, and AIPP3-FLAG over Pol V peaks (n=10,868). ChIP-seq signal of control samples were subtracted for plotting. **c**, Screenshots of Pol V, MOM1-Myc, PIAL2-Myc, AIPP3-FLAG and PHD1-FLAG ChIP-seq signals with control ChIP-seq signal subtracted and CG, CHG, and CHH DNA methylation level by WGBS over representative RdDM sites. **d**, Volcano plot showing proteins that have significant interactions with MOM1 as detected by crosslinking IP-MS, with RdDM pathway components and MOM1 complex components labeled. Crosslinking IP-MS of Col-0 plant tissue was used as control. **e**, AIPP3-FLAG ChIP-seq peaks were divided into two groups: Group 1 peaks (n = 3075) have MOM1-Myc ChIP-seq signal enriched and Group 2 peaks (n = 523) have no enrichment of MOM1-Myc ChIP-seq signal. Metaplots and heatmaps representing ChIP-seq signals of MOM1-Myc, AIPP3-FLAG, PHD1-FLAG and PHD3-FLAG over these two groups of AIPP3 peaks are shown. ChIP-seq signal of control samples were subtracted for plotting.

To study the function of the MOM1 complex, ChIP-seq was performed in FLAG or MYC tagged MOM1, PIAL2, PHD1 and AIPP3 transgenic lines. Surprisingly, MOM1, PHD1, AIPP3, and PIAL2 were all highly colocalized with Pol V at RdDM sites (Fig. 1 b and c). To further validate colocalization of the MOM1 complex with the RdDM sites, we performed crosslinking IP-MS of FLAG tagged MOM1 and observed that in addition to the MOM1 complex components, several proteins in the RdDM machinery, including NRPD2 (subunit of Pol-V and Pol-IV), NRPE1 (subunit of Pol-V), DMS3 and SPT5L (P=0.01243) were also significantly enriched (Fig. 1d and Supplementary Table 2). Interestingly, we also observed a significant enrichment of MORC1 and MORC6 in the MOM1 crosslinking IP-MS (Fig. 1d and Supplementary Table 2), suggesting that the RdDM machinery, the MORC proteins and the MOM1 complex are co-located at the same loci, either because they are crosslinked by co-bound stretches of chromatin, or because the crosslinking process enhanced relatively weak interactions between the proteins.

Further examination of the MOM1 ChIP-seq signal over the AIPP3 peaks suggested that a group of AIPP3 binding loci were not enriched for MOM1 (Fig. 1e). We named the group of AIPP3 peaks that have MOM1 ChIP-seq signal enriched as Group 1 peaks and those with no MOM1 enrichment as Group 2 peaks. Consistent with our IP-MS data suggesting that PHD1 is a MOM1 complex component, PHD1 ChIP-seq signal was predominantly enriched in Group1 AIPP3 peaks which also bound to MOM1 (Fig. 1e and Supplementary Fig1). We also performed Chip-seq with FLAG tagged PHD3 transgenic plants. In contrast to PHD1, PHD3 ChIP-seq signal was enriched in both groups of AIPP3 peaks, closely resembling the pattern of AIPP3 ChIP-seq signal (Fig. 1e and Supplementary Fig1). These data further suggests that AIPP3 exists in multiple protein complexes including the MOM1 complex.

### Zinc finger tethering of MOM1 complex components to the *FWA* promoter triggers DNA methylation and silencing

Since MOM1 is localized to RdDM sites, and ZF fusions of RdDM components have been shown to silence *FWA* expression in the *fwa* mutant^43^, we investigated whether tethering the components of the MOM1 complex could also lead to the silencing of *FWA* expression. We created ZF fusion proteins with MOM1, MOM2, PIAL1, PIAL2, AIPP3 and PHD1 and transformed them into the *fwa* mutant. ZF fusion of MOM1, MOM2, PIAL1, PIAL2 and PHD1 restored the early flowering phenotype (Fig. 2a, Supplementary Fig. 2a), significantly repressed *FWA* expression (Fig. 2b), and induced DNA methylation at the *FWA* promoter region as detected by the bisulfite amplicon sequencing analysis (BS-PCR-seq) (Fig. 2c). The DNA methylation induced at the *FWA* promoter region was retained in the transgene-free T2 plants, showing that the newly established DNA methylation was heritable (Fig. 2c). PIAL1-ZF was somewhat less efficient at restoring the early flowering phenotype in the T1 population (Supplementary Fig.2a). However, reduced *FWA* mRNA levels and increased *FWA* promoter DNA methylation, as measured with McrBC digestion assay, were detected in some PIAL1-ZF T1 plants (Supplementary Fig. 2b), and plants with similar flowering time to the Col-0 were observed from the three T2 populations of the earliest flowering T1 plants (Fig. 2a, Supplementary Fig.2a). In addition, DNA methylation at the *FWA* promoter region retained in T2 plants free of PIAL1-ZF transgenes, showing that PIAL1-ZF can also induce heritable DNA methylation (Fig. 2c). AIPP3-ZF led to a slightly early flowering time in the T1 population compared to the *fwa* control population, however, zero T1 transgenic plants and very few T2 plants flowered as early as the Col-0 control plants (Supplementary Fig 2a and c). A low level of DNA methylation in the *FWA* promoter region, mainly methylation in the CHH sequence context, was detected in the AIPP3-ZF T2 plants which were positive for the transgene (Supplementary Fig. 2d). However, no DNA methylation was detected in transgene-free T2 plants segregating in the same T2 populations (Supplementary Fig. 2d). These data suggests that the establishment of DNA methylation by AIPP3-ZF is much weaker compared to other MOM1 complex components. Previous work reported that, in addition to the designed binding site in the *FWA* promoter, ZF also binds to many off-target sites in the genome^43^. Whole genome bisulfite sequencing (WGBS) showed that MOM1-ZF, MOM2-ZF, PIAL1-ZF, PIAL2-ZF and PHD1-ZF also enhanced DNA methylation at ZF off-target sites (Fig. 2d and Supplementary Fig. 3a). Overall, these results suggest that ZF fusions of the components of the MOM1 complex are able to trigger the establishment of DNA methylation and silence *FWA* expression in the *fwa* mutant, as well as establish methylation at other ZF off target sites.

**Fig. 2.**
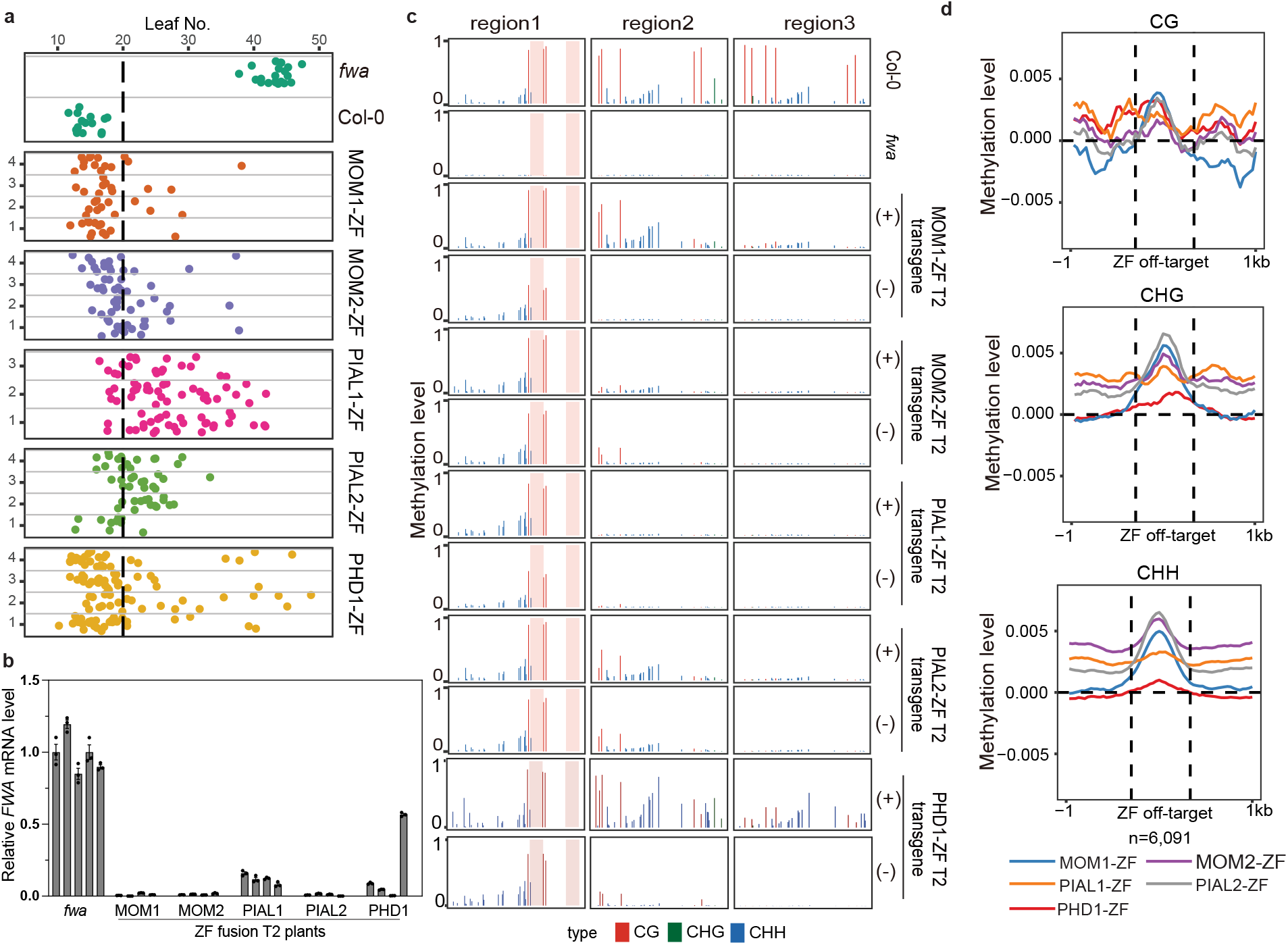
ZF tethering of the MOM1 complex to the *FWA* promoter triggers DNA methylation *FWA* silencing. **a**, Flowering time of *fwa*, Col-0 and representative T2 lines of MOM1-ZF, MOM2-ZF, PIAL1-ZF, PIAL2-ZF and PHD1-ZF in the *fwa* background. The numbers of independent plants (n) scored for each population are listed in Supplementary Table 5. **b**, qRT-PCR showing the relative mRNA level of *FWA* gene in the leaves of *fwa* plants, and four T2 plants of MOM1-ZF, MOM2-ZF, PIAL1-ZF, PIAL2-ZF and PHD1-ZF in the *fwa* background. Bar plots and error bars indicate the mean and standard error of three technical replicates, respectively, with individual technical replicates shown as dots. **c**, CG, CHG, and CHH DNA methylation levels over *FWA* promoter regions measured by BS-PCR-seq in Col-0, *fwa* and representative T2 plants of MOM1-ZF, MOM2-ZF, PIAL1-ZF, PIAL2-ZF and PHD1-ZF in the *fwa* background with (+) or without (-) corresponding transgenes. Pink vertical boxes indicate ZF binding sites. **d**, Metaplots showing relative variations (sample minus control) of CG, CHG, and CHH DNA methylation levels over ZF off-target sites in representative T2 plants of MOM1-ZF, MOM2-ZF, PIAL1-ZF, PIAL2-ZF and PHD1-ZF in the *fwa* background versus *fwa* control plants measured by whole genome bisulfite sequencing (WGBS).

The CMM2 domain has been shown to essential for the transcriptional gene silencing function of the MOM1 protein^12,13^. We found that a ZF fusion with the CMM2 domain together with a nuclear localization signal (called miniMOM1)^12^ was efficient at targeting heritable *FWA* methylation (Supplementary Fig. 3b and c). We performed IP-MS with a miniMOM1-FLAG line and found peptides for MOM2, PIAL1, and PIAL2, but not for AIPP3 or PHD1 (Supplementary Table 1). These results suggest that AIPP3 and PHD1 may be dispensable for the targeting of methylation to *FWA* promoter.

To begin to dissect the requirements for MOM1-mediated establishment of *FWA* methylation and silencing, we first transformed MOM1-ZF and PHD1-ZF into *mom1 fwa* and *phd1 fwa* mutant backgrounds (Supplementary Fig. 4a). MOM1-ZF was able to trigger early flowering in *phd1 fwa*, positioning MOM1 downstream of PHD1 (Supplementary Fig. 4a). Consistent with this order of action, the *mom1* mutant blocked PHD1-ZF activity (Supplementary Fig. 4a). PHD1-ZF activity was also blocked in the *aipp3 fwa* mutant (Supplementary Fig. 4a). These results are consistent with IP-MS result showing that the MOM1-PHD1 interaction was abolished in the *aipp3-1* mutant (Supplementary Table 1).

To further dissect the hierarchy of action of MOM1 components, we transformed PIAL2-ZF into *aipp3 fwa, phd1 fwa, mom2 fwa* and *mom1 fwa* mutant backgrounds and found that PIAL2-ZF triggered an early flowering phenotype in all mutant backgrounds (Supplementary Fig. 4b), suggesting that PIAL2 might act at the most downstream position within the MOM1 complex. However, we also transformed MOM1-ZF into *aipp3 fwa, mom2 fwa* and *pial1/2 fwa*, and found that MOM1-ZF was also able to trigger early flowering in all these mutant backgrounds (Supplementary Fig. 4a), suggesting that MOM1 acts at a step parallel with PIAL1/2 in targeting DNA methylation. We did however observe that MOM1-ZF showed a lower efficiency of triggering early flowering in the *pial1/2 fwa* mutant compared to wild type or the other mutants (Supplementary Fig. 2a and 4a), suggesting that PIAL1/2 is required for the full functionality of MOM1-ZF. We also transformed MOM2-ZF into *aipp3 fwa, phd1 fwa, mom1 fwa*, and *pial1/2 fwa*, and like MOM1-ZF and PIAL2-ZF, MOM2-ZF was able to trigger early flowering in all the mutants (although again with lower efficiency in the *pial1/2 fwa* background) (Supplementary Fig. 4b), suggesting that MOM2 also acts with MOM1 and PIAL2 in a very downstream step in triggering methylation, but that PIAL1/2 is required for its full functionality. As a control, we compared the flowering time in the mutant backgrounds without transgenes. *mom1 fwa* flowers at similar time compare to *fwa*, while *pial1/2 fwa* and *aipp3 fwa* flowered slightly earlier (Supplementary Fig. 4c), suggesting that the deficiency in triggering early flowering by ZF fusion proteins in these backgrounds is not due to differences in flowering time of mutant backgrounds. In summary, these results suggest that MOM1, PIAL1/PIAL2, and MOM2 are acting as the most downstream factors in the MOM1 complex for establishing DNA methylation at the *FWA* promoter.

### MOM1-ZF recruits the Pol V arm of the RdDM machinery via MORC6 to establish *de novo* DNA methylation at the *FWA* promoter

Because the tethering of RdDM components to *FWA* has been previously shown to efficiently establish methylation of *FWA*^28,43^, we hypothesized that MOM1-ZF established *FWA* DNA methylation by recruiting the RdDM machinery. To test this hypothesis, we transformed PIAL2-ZF and MOM1-ZF into *fwa* backgrounds in which RdDM mutations had been introgressed, including *nrpd1, suvh2/9, dms3, drd1, rdm1, nrpe1*, and *drm1/2*^43^. PIAL2-ZF and MOM1-ZF were still capable of triggering an early flowering phenotype in *nrpd1* (the largest subunit of Pol IV), suggesting that siRNA biogenesis was not needed for methylation targeting (Fig. 3a). These fusions were also capable of triggering silencing in the *suvh2/9* mutant background (Fig. 3a), showing that the SUVH2 and SUVH9 factors that normally recruit the DDR complex and Pol V to chromatin were not needed for silencing. However, silencing activity of PIAL2-ZF and MOM1-ZF was blocked by DDR component mutations (*dms3, drd1*, and *rdm1*) as well as by mutations in the largest subunit of Pol V (*nrpe1*) and the DRM de novo methyltransferases (*drm1/2*) (Fig. 3a). These results place the action of PIAL2-ZF and MOM1-ZF upstream of the DDR complex. Interestingly, it was previously shown that, MORC6-ZF showed an identical pattern of triggering *FWA* methylation in wild type, *nrpd1*, and *suvh2/9*, but not in *dms3, drd1, rdm1, nrpe1*, or *drm1/2*^43^. This similarity prompted us to test the targeting of PIAL2-ZF, MOM1-ZF, MOM2-ZF, PIAL1-ZF and PHD1-ZF in the *morc6 fwa* genetic background. Interestingly, we found that all these ZF fusions failed to trigger *FWA* silencing in *morc6* (Fig. 3a and Supplementary Fig. 4d), suggesting that the MOM1 complex acts upstream of MORC6. To further confirm this order of action we transformed MORC6-ZF into *fwa* backgrounds in which the *mom1-3, mom2-1, pial1/2, phd1-2* and *aipp3-1* mutants had been introgressed. We found that MORC6-ZF could successfully target silencing of *FWA* in all these backgrounds (Supplementary Fig. 4d), confirming that MORC6 acts downstream of the MOM1 complex in the targeting of *FWA* silencing. We also performed ChIP-seq of MYC-tagged MORC6 in the *morc6-3* mutant background. Similar to the MOM1 complex reported here, and similar to that previously reported for MORC4 and MORC7 proteins^40^, we observed that MORC6 was highly colocalized with Pol V at RdDM sites (Fig. 3b and c).

**Fig. 3.**
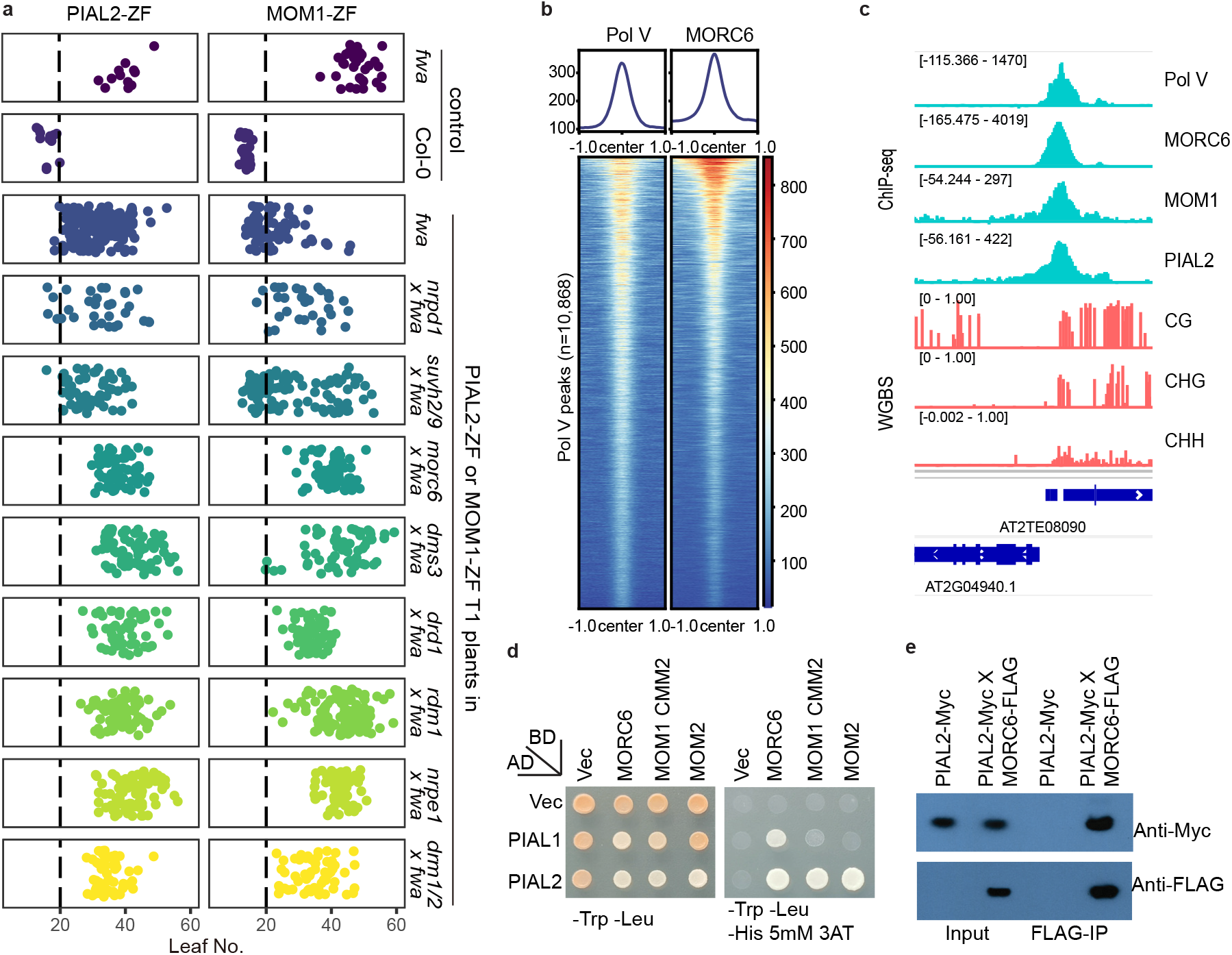
MOM1-ZF recruits the Pol V arm of the RdDM machinery via MORC6. **a**, Flowering time of *fwa*, Col-0, and T1 lines of PIAL2-ZF and MOM1-ZF in the *fwa* mutant backgrounds as well as in backgrounds of *fwa* introgressed mutants, including *nrpd1, suvh2/9, morc6, dms3, drd1, rdm1, nrpe1* and *drm1/2*. The numbers of independent plants (n) scored for each population are listed in Supplementary Table 5. **b**, Metaplots and heatmaps representing ChIP-seq signals of Pol V and MORC6-Myc over Pol V peaks (n=10,868). ChIP-seq signal of control samples were subtracted for plotting. **c**, Screenshots of Pol V, MORC6-Myc, MOM1-Myc and PIAL2-Myc ChIP-seq signals with control ChIP-seq signals subtracted and CG, CHG, and CHH DNA methylation level by WGBS over a representative RdDM site. **d**, Yeast Two-Hybrid assay showing *in vitro* direct interactions between PIAL1 and PIAL2 with MORC6 and the MOM1 CMM2 domain, as well as between PIAL2 and MOM2. **e**, PIAL2 and MORC6 *in vivo* interaction shown by co-immunoprecipitation (Co-IP) in MORC6-FLAG and PIAL2-Myc crossed lines.

Given that PIAL1/PIAL2, MOM1, and MOM2 appeared to be the most downstream critical components of the MOM1 complex required for triggering *FWA* methylation, and that ZF fusions of these proteins failed to trigger methylation in a *morc6* mutant, we reasoned at least one of these components might physically interact with MORC6. Indeed, we found that PIAL2 was able to interact with MORC6 in a Yeast Two-Hybrid assay (Fig. 3d). We also confirmed this interaction by an *in vivo* co-immunoprecipitation assay, observing that MORC6-FLAG was able to interact with PIAL2-Myc (Fig. 3e). While there could certainly be other important interactions, these results suggest that the MOM1 complex likely recruits MORC6 in part via a physical interaction between PIAL2 and MORC6. MORC6 then triggers *FWA* methylation via its interaction with the RdDM machinery as previously reported^40^.

### The MOM1 complex facilitates the process of transgene silencing

Several previous screens identified MOM1 as a key component in the maintenance of the silenced state of the transgene reporters used in the screen^3,5,47^. RdDM is involved in the maintenance of DNA methylation, but also in the initial establishment of methylation. For example, studies have shown that when an extra copy of the *FWA* gene is introduced into *Arabidopsis* plants via Agrobacterium-mediated transformation, it is very efficiently methylated and silenced in the wild type background. However, this methylation and silencing is blocked in RdDM mutants, leading to overexpression and a late flowering phenotype^15,29,48^. Interestingly, the silencing of *FWA* transgenes was previously shown to be less efficient in the *morc* mutants^40^. Since the MOM1 complex is closely linked with the RdDM machinery and MORC6, we suspected that the MOM1 complex may also facilitate the efficient establishment of transgene silencing. To test this, the *FWA* transgene was transformed into Col-0 plants (wild type) and the mutant background of *nrpe1-11, mom1-3, pial1/2, mom2-22, aipp3-1* and *phd1-2*. As expected^40^, the T1 transgenic plants in the *nrpe1-11* background flowered much later (mean leaf number: 33.81) compared to those in the Col-0 background (mean leaf number: 15.91) (Fig. 4a and Supplementary Fig. 5a). We found that T1 plants containing the *FWA* transgene in *mom1-3* or *pial1/2* mutant backgrounds also flowered later than in those in the Col-0 background, with a mean leaf number of 27.55 (*mom1)* and 31.98 (*pial1/2)* (Fig. 4a and Supplementary Fig. 5a). We examined four late flowering T1 plants in each of the *mom1-3* and *pial1/2* mutant backgrounds and observed that, consistent with their late flowering phenotype, *FWA* mRNA levels were higher than in the Col-0 background (Fig. 4b upper panel). The unmethylated *FWA* promoter DNA fraction, as detected with McrBC digestion assay, was also higher in these T1 plants compared to Col-0, suggesting that efficient establishment of DNA methylation on the *FWA* transgene is impaired in *mom1-3* and *pial1/2* mutants (Fig. 4b lower panel).

**Fig. 4.**
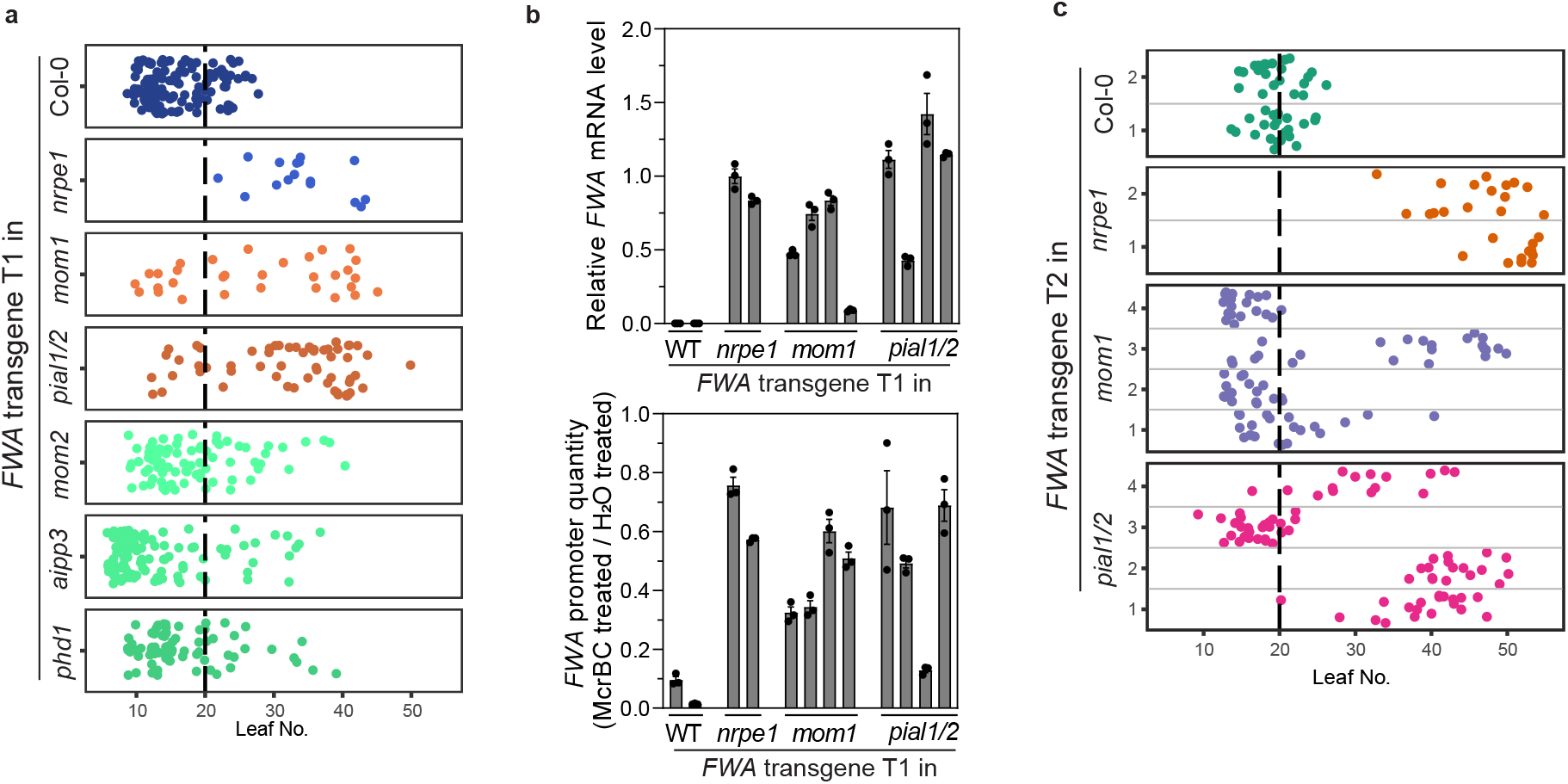
The MOM1 complex facilitates the process of transgene silencing. **a**, Flowering time of *FWA* transgene T1 plants in the Col-0, *nrpe1-11, mom1-3, pial1/2, mom2-2, aipp3-1* and *phd1-2* genetic backgrounds. **b**, Relative *FWA* mRNA level (upper panel) and relative *FWA* promoter DNA quantity after McrBC treatment (lower panel) of four late-flowering *FWA* transgene containing T1 plants in the *mom1-3* and *pial1/2* genetic backgrounds. *FWA* transgene containing T1 plants in the Col-0 and *nrpe1-11* backgrounds were used as controls. Bar plots and error bars indicate the mean and standard error of three technical replicates, respectively, with individual technical replicates shown as dots. **c**, Flowering time (leaf number) of *FWA* transgene T2 plants in the Col-0, *nrpe1-11, mom1-3* and *pial1/2* genetic backgrounds. For **a** and **c**, the numbers of independent plants (n) scored for each population are listed in Supplementary Table 5.

Although a small number of T1 *FWA* transgenic plants with a late flowering time was also observed in the *mom2-2, aipp3-1* and *phd1-2* backgrounds, the average flowering time of these T1 plants was not significantly later than that of the T1 plants in the Col-0 background (Fig. 4a and Supplementary Fig. 5a). In fact, the *FWA* transgene T1 population in the *aipp3-1* background flowered significantly earlier than in Col-0 (Supplementary Fig. 5a), likely due to the fact that the *aipp3-1* mutant itself flowers earlier than Col-0 plants (Supplementary Fig. 5b), as previously reported^46^. These data suggests that MOM2, AIPP3 and PHD1 contribute minimally to efficient silencing of the *FWA* transgene, whereas MOM1 and PIAL1/2 contribute significantly.

In strong RdDM mutants such as *nrpe1*, the *FWA* transgene stays unmethylated and all of the T2 offspring plants with the *FWA* transgene show a late flowering phenotype^40^. We grew the T2 populations of four late flowering T1 plants in each of the *mom1-3* and *pial1/2* backgrounds and scored for their flowering time. In T2 plant populations in *mom1-3* line 2 and line 4, as well as in *pial1/2* line 3, all transgene positive plants showed a relatively early flowering time, similar to controls of T2 plants with *FWA* transgene in Col-0 background (Fig. 4c). However, in the other T2 populations tested, we observed transgene positive plants with flowering time spanning from very late to early (*mom1-3* T2 line 1 and line3, in *pial1/2* T2 line 1 and line 4), as well as one line with 100% late flowering plants (*FWA* transgene in *pial1/2* line 2) (Fig. 4c). These data suggests that instead of completely blocking *FWA* transgene silencing as in strong RdDM mutants like *nrpe1*, mutation of *MOM1* or *PIAL1/2* reduces the efficiency of *FWA* transgene silencing, similar to what was previously observed for mutation of *MORC* genes^40^.

### The MOM1 complex influences DNA methylation and chromatin accessibility at some endogenous RdDM sites

The strong co-localization of the MOM1 complex with RdDM sites suggests that the MOM1 complex might facilitate the endogenous function of the RdDM machinery. To test this hypothesis, we performed Whole Genome Bisulfite Sequencing (WGBS) in *phd1-2, phd1-3, aipp3-1*, and *mom2-2* and analyzed these together with previously published WGBS data from the *morc6-3*^24^, *morc1/2/4/5/6/7 hextuple* (*morchex)*^39^, *mom1-3* and *pial1/2* mutants^5^, followed by analysis using the High-Confidence Differentially Methylated Regions (hcDMRs) pipeline^8^. We observed a little over 200 hypo CHH hcDMRs in *mom1-3*and *pial1/2* double mutant and 23 hypo CHH hcDMRs in *mom2-2*, most of which overlapped with those of *morc6* and *morchex* at RdDM sites (520 DMRs in morchex)^39^ (Figure 5a and 5b, Supplementary Table 3). This is consistent with an earlier analysis that showed a strong overlap of *mom1* hypomethylated DMRs with those of the *morchex* mutant^8^. On the other hand, the *aipp3-1* mutant only shared 1 out of its 13 hypo CHH hcDMRs with *morc6* (Supplementary Table 3), and neither of the *phd1* mutant alleles tested showed any hypo CHH hcDMRs (Supplementary Table 3). To further explore the functions of MOM1 complex components at these sites, we performed RNA-seq in Col-0, *morc6-3, morchex*^39^ and mutants of the MOM1 complex components. We observed that expression level of the genomic regions within 1 kb of the 520 CHH hypo-DMR regions previously found in the *morchex* mutant were slightly upregulated in *mom1-3, pial1/2, morc6-3* and *morchex* mutants, but not in *phd1-2, aipp3-1, pial1-2, pial2-1*, or *mom2-2* mutants (Figure 5c), showing that MOM1/PIAL1/PIAL2, along with MORCs, are required for the maintenance of CHH methylation and gene silencing at a small subset of RdDM sites, while AIPP3, PHD1, and MOM2 seem to play little role in this process.

**Fig. 5.**
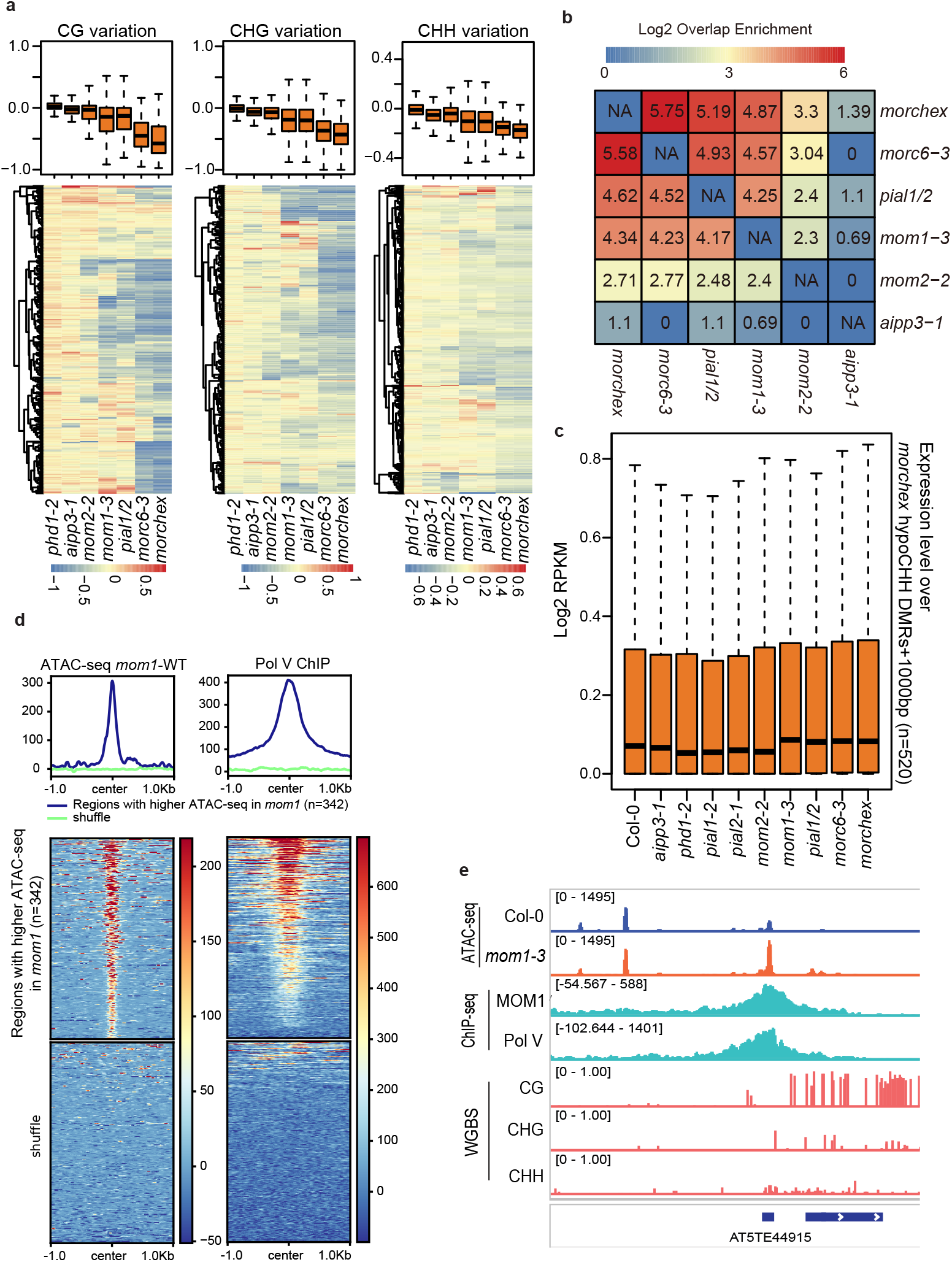
The MOM1 complex influences DNA methylation and chromatin accessibility at some endogenous RdDM sites. **a**, Boxplots and heatmaps showing the variation of CG, CHG, and CHH DNA methylation in *phd1-2, aipp3-1, mom2-2, mom1-3, pial1/2, morc6-3* and *morchex* mutants vs Col-0 wild type over hypo CHH hcDMRs of the *morchex* mutant (n=520). **b**, Heatmap depicting the overlapping enrichment of hypo CHH hcDMRs among *aipp3-1, mom2-2, mom1-3, pial1/2, morc6-3* and *morchex* mutants over *morchex* mutant hypo CHH hcDMRs (n=520). **c**, Boxplot representing the expression level (RNA-seq signal normalized by RPKM) of the genomic bins of 1 kb from hypo CHH hcDMRs (n=520) of the *morchex* mutant in Col-0, *aipp3-1, phd1-2, pial1-2, pial2-1, mom2-2, mom1-3, pial1/2, morc6-3* and *morchex* mutants. **d**, Metaplots and heatmaps representing ATAC-seq signal (*mom1-3* minus Col-0) and Pol V ChIP-seq signal (subtracting control ChIP-seq signal) over regions with higher ATAC-seq signals in *mom1-3* (n=342) and shuffled regions. **e**, Screenshots of ATAC-seq signals of Col-0 and *mom1-3*, ChIP-seq signals of MOM1-Myc and Pol V (subtracting control signal) as well as CG, CHG, and CHH DNA methylation level by WGBS over a representative RdDM site. In box plots of **a** and **c**, center line represents the median; box limits represent the 25th and 75th percentiles; whiskers represent the minimum and the maximum.

We also performed ATAC-seq and detected 342 regions with increased ATAC-seq signal in the *mom1-3* mutant compared to Col-0 (Fig 5d). We also found that Pol V Chip-seq signal was highly enriched in these regions (Fig 5d and 5e), suggesting that the MOM1 complex reduces chromatin accessibility at a subset of RdDM sites. Together, these results suggest that the MOM1 complex contributes to the endogenous function of the RdDM machinery, facilitating the maintenance of DNA methylation and a more closed chromatin status at some RdDM sites.

### The MOM1 complex has endogenous function divergent from the RdDM machinery

Previous studies have shown that the *mom1* mutants show derepression of pericentromeric heterochromatin regions, while the targets of the RdDM machinery tends to locate in euchromatic regions of the chromosome arms^5,6,49,50^. Consistent with these differences, we observed that ChIP-seq signals of MOM1, MORCs, and to a lesser extent PIAL2 were more highly enriched on transposable elements (TEs) located in pericentromeric regions as compared to TEs located in the chromosome arms – the opposite pattern to that of Pol V ChIP-seq^34^ (Figure 6a). From our RNA-seq, *mom1* and *pial1/2* mutants also showed transcriptional upregulation mainly in pericentromeric regions, while up-regulated TEs in the *nrpe1-11* mutant were located more broadly over the chromosomes including both pericentromeric regions and the euchromatic arms (Supplementary Fig 6a). Consistent with previous reports^7^, *morc6-3* and *morchex* mutants also displayed derepression of pericentromeric regions (Supplementary Fig 6a). Upregulated differentially expressed TEs (DE-TEs) in the *morc6-3* and *morchex* mutants^39^ showed a prominent overlap with those of the *mom1-2, mom1-3*, and *pial1/2* mutants (Supplementary Fig 6b). The *phd1, aipp3*, and *mom2* mutants on the other hand showed little change in expression at these same sites (Supplementary Fig 6a and 6b)., suggesting that these factors are less important for this silencing function.

**Fig. 6.**
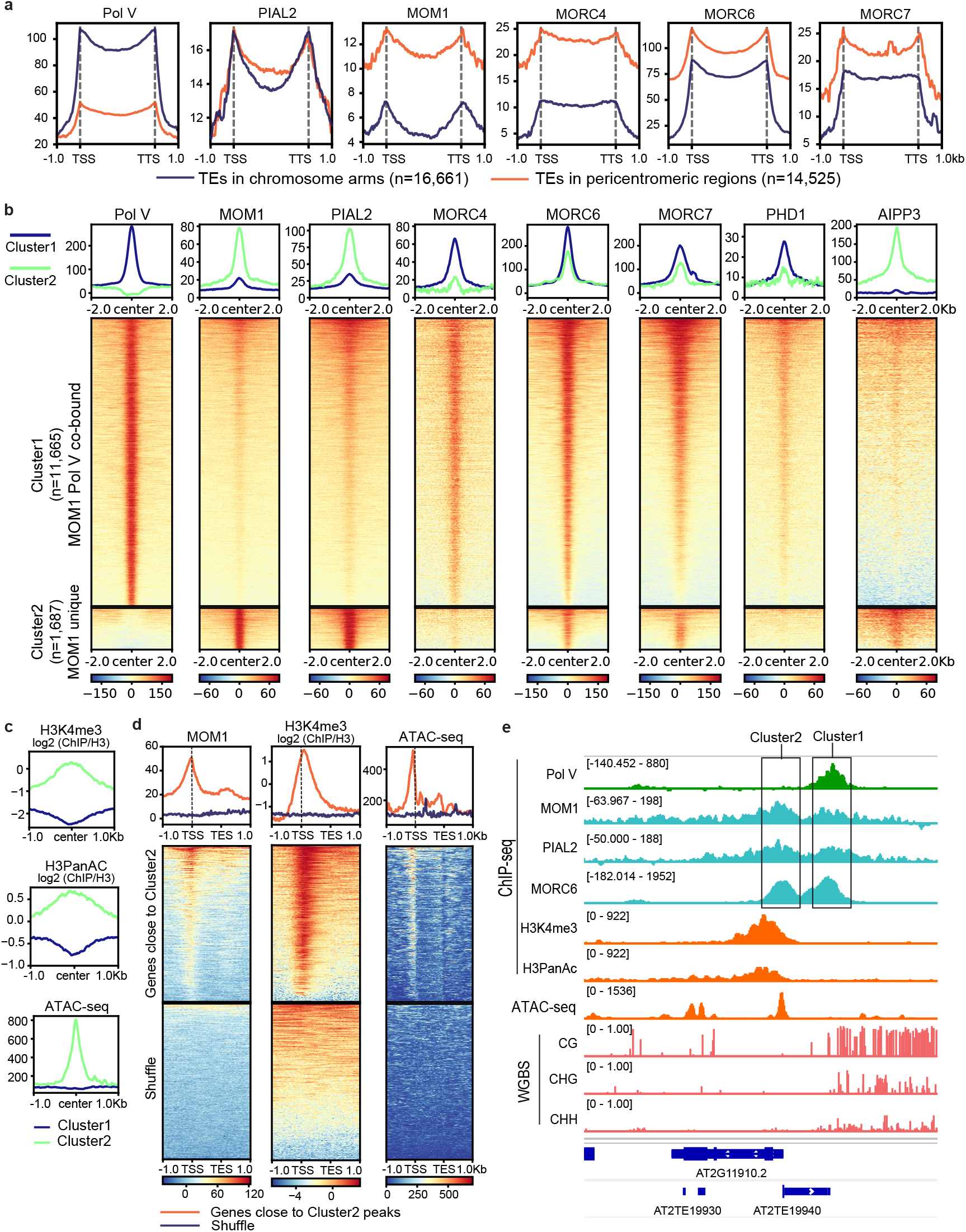
MOM1 complex components and MORCs shows genomic distribution patterns distinct from that of the RdDM component Pol V. **a**, Metaplots of ChIP-seq signals of Pol V, PIAL2, MOM1, MORC4, MORC6, and MORC7 over TEs in euchromatic arms (n=16,661) and TEs in pericentromeric regions (n=14,525), with control ChIP-seq signals subtracted. **b**, Metaplots and heatmaps of ChIP-seq signals of Pol V, MOM1, PIAL2, MORC4, MORC6, MORC7, PHD1, and AIPP3 over Cluster 1 and Cluster 2 ChIP-seq peaks of MOM1 and Pol V, with control ChIP-seq signals subtracted. **c**, Metaplots of ChIP-seq signals of H3K4me3 and H3PanAC (normalized to H3), as well as ATAC-seq signal of Col-0 over Cluster 1 and Cluster 2 peaks of MOM1 and Pol V. **d**, Metaplots and heatmaps of MOM1 ChIP-seq signal (with control ChIP-seq signal subtracted), H3K4me3 ChIP-seq signal (normalized to H3) and ATAC-seq signal of Col-0 plants over genes close to Cluster 2 peaks and shuffled control regions. **e**, Screenshots of Pol V, MOM1, PIAL2, MORC6 ChIP-seq signals with control ChIP-seq signals subtracted, H3K4me3 and H3PanAC ChIP-seq signals, ATAC-seq signal of Col-0 plants, as well as CG, CHG, and CHH DNA methylation level by WGBS over a representative genomic region containing both Cluster 1 and Cluster 2 ChIP-seq peaks.

We also discovered a set of MOM1 ChIP-seq peaks that did not overlap with DNA methylation. We initially discovered these by performing unsupervised clustering of MOM1 ChIP-seq data with Pol V ChIP-seq data^34^, and identified a group of MOM1 unique peaks not colocalizing with Pol V sites (Fig 6b). We named the MOM1 and Pol V co-binding peaks as Cluster 1 peaks and the MOM1 unique peaks as Cluster 2 peaks (Fig 6b). Other components of the MOM1 complex, such as the PIAL2, AIPP3 and to a lesser extent, PHD1 were also enriched at cluster 2 peaks (Fig 6b). In addition, MORC4^40^, MORC6 and MORC7^40^ co-localized with MOM1 at both the RdDM sites and the MOM1 unique Cluster 2 peaks (Fig 6b). Interestingly, we found that the Cluster 2 peaks were enriched for active histone marks H3K4me3 and H3PanAc^51^, as well as accessible chromatin indicated by ATAC-seq signal (Fig 6c). This observation is consistent with a recent study reporting that MORC7 protein binds to active chromatin regions devoid of RdDM^40^. While H3K4me3 tends to peak after the Transcription Start Site (TSS), the MOM1 ChIP-seq signal tended to peak around the TSS of the genes near Cluster 2 peaks, similar to the ATAC-seq signal (Fig 6d and 6e). The function of the MOM1 complex at these non-DNA methylated sites is currently unknown.

Overall, the ChIP-seq data suggests that while MOM1 and PIAL2 show strong localization to RdDM sites, they and the MORC proteins are more enriched in pericentromeric regions compared to the RdDM machinery. In addition, they are also present at unique active chromatin sites. The recruitment mechanism and the endogenous function of the MOM1 complex binding at the active chromatin sites need to be further investigated.

## Discussion

Due to the lack of major change in DNA methylation status in derepressed transgenes and endogenous TEs in the *mom1* mutant, MOM1 function has long been considered as independent of DNA methylation or downstream of DNA methylation. In our study, we observed a close link between the MOM1 complex and the RdDM machinery. By tethering the MOM1 complex with ZF in the *fwa* mutant, heritable DNA methylation was established at the *FWA* promoter, suggesting that the RdDM machinery was recruited as a result. Consistent with this, silencing and methylation of *FWA* were blocked in mutants of the DDR complex, as well as the *nrpe1* and *drm1/2* mutants, but not in the *suvh2/9* and *nrpd1* mutants. Thus, the recruitment of the DRM2 *de novo* DNA methyltransferase by the MOM1 complex requires the Pol V arm of the RdDM pathway. Previous MORC6-ZF tethering experiments resulted in similar results, *i*.*e*., the DDR complex and the downstream Pol V arm was required for silencing of *FWA*. In addition, we found that mutation of *MORC6* blocked *FWA* silencing mediated by ZF fusion to MOM1 complex components, suggesting that the MOM1 complex recruits the RdDM machinery via MORC6. This was also consistent with our observed physical interaction between PIAL2 of the MOM1 complex and MORC6. These observations do not however exclude the possibility that physical interactions might also exist between MOM1 complex components and other components of the RdDM machinery.

We also found that MOM1 and PIAL1/2 are required for the efficiency of the establishment of methylation and silencing of *FWA* transgenes. Compared to RdDM mutants that completely block DNA methylation and silencing of *FWA* transgenes, the *mom1* and *pial1/2* mutants only showed a reduced efficiency of silencing, similar to what was observed in the *morchex* mutant. How the MOM1 complex performs this function is unclear. The MOM1 complex might facilitate the initial loading of the RdDM machinery onto the *FWA* transgene, or it might allow for greater retention of the loaded RdDM machinery for more efficient DNA methylation and silencing, as has been suggested for the MORCs^40^. It is also possible that MOM1 complex mutants show defective transcriptional silencing of *FWA* during the DNA methylation establishment process, such that positive epigenetic marks associated with transcription may compete with the methylation establishment process, making it slower or less efficient. Consistent with the connections between MOM1 and RdDM revealed by ZF tethering results and *FWA* transgene silencing results, our ChIP-seq data showed that the MOM1 complex highly co-localized with RdDM sites in the genome. Our analysis of WGBS data also showed that *MOM1* and *PIAL1/2* were required to maintain CHH methylation at a small subset of RdDM sites, which significantly overlap with CHH hypoDMR sites in the *morchex* mutants. A previous study also reported a similar observation with WGBS data from a different *mom1* mutant allele (*mom1-2*)^8^. Thus, aside from the previous findings that that transgene and TE silencing are released in the *mom1* mutant background without major DNA methylation changes^3,5,7^, the MOM1 complex^8^, together with the MORC proteins, are also required for the maintenance of DNA methylation at a small subset of RdDM sites. It seems likely that this would be mechanistically related to the role of both MOM1 and MORCs in the establishment of *FWA* transgene silencing, and it is intriguing to speculate that this might reflect an ancient role of these proteins in the initial establishment of methylation and silencing of novel invading transposable elements.

The role of MOM1 in establishment and maintenance of RdDM described in this study is clearly not the only role of MOM1 in epigenome regulation since comparison of DE-TEs and DE-genes in the *nrpe1* and *mom1* mutants in previous studies^5,6^ indicates that the majority of their endogenous targets do not overlap. In addition, some genes are specifically upregulated in the *mom1 nrpe1* double mutant showing that MOM1 and RdDM clearly have some non-overlapping functions^6^. The localization of the MOM1 complex at RdDM sites might be needed for the repression of these common target sites upregulated in *mom1 nrpe1*. In addition, we observed that the MOM1 complex, as well as the MORC proteins, showed a stronger enrichment over TEs in the pericentromeric region, a pattern that is the opposite of Pol V, which shows stronger enrichment over TEs in the euchromatic arms. Thus, although the MOM1 complex and MORC proteins broadly co-localizes with most RdDM sites, they have different binding profiles compared to the core RdDM component Pol V.

In addition to the localization at RdDM sites, we identified a unique set of MOM1 peaks which are enriched with active chromatin marks. This is reminiscent of an earlier study reporting that MOM1 regulates transcription in intermediate heterochromatin, which is associated with both active and repressive histone marks^49^. Interestingly, the MOM1 complex and MORCs seem to behave similarly in binding active chromatin, as MORC7 was also reported to bind active chromatin devoid of RdDM^52^, and MORCs are colocalized at these MOM1 unique peaks. The mechanism of recruiting the MOM1 complex to these unique peaks and the function of MOM1 at these active chromatin sites is unknown.

In summary, our results uncover a new function for the MOM1 complex in the efficiency of both the establishment and maintenance of RNA-directed DNA methylation and gene silencing, and point to a potential function at some unmethylated euchromatic regions, suggesting that MOM1 plays multifaceted roles in epigenome regulation.

## Materials and Methods

### Growth condition, molecular cloning and plant materials

*Arabidopsis thaliana* plants in this study were Col-0 ecotype and were grown under 16h light: 8h dark condition. The T-DNA insertion lines used in this study are: *aipp3-1* (GABI_058D11), *aipp3-2* (SAIL_1246_E10), *mom1-2* (SAIL_610_G01), *mom1-3* (SALK_141293), *mom1-7* (GABI_815G11), *mom2-1* (WiscDsLox364H07), *mom2-2* (SAIL_548_H02), *pial1-2* (CS358389), *pial2-1* (SALK_043892), *morc6-3* (GABI_599B06), *aipp2-1* (SALK_057771), *nrpe1-11* (SALK_029919) and *morchex*^39^ consisting of *morc1-2* (SAIL_893_B06), *morc2-1* (SALK_072774C), *morc4-1* (GK-249F08), *morc5-1* (SALK_049050C), *morc6-3* (GABI_599B06), and *morc7-1* (SALK_051729). In addition to the T-DNA insertion line, three *phd1* mutant alleles were generated using a YAO promoter driven CRISPR/Cas9 system^53^. *phd1-2* contained a single nucleotide T insertion and *phd1-3* contained a 13-nucleotide deletion and an 18-nucleotide duplication in the 2^nd^ exon of PHD1 gene, both of which led to early termination of the protein at amino acid 53 located within the PHD domain. *phd1-4* contained a single nucleotide T insertion in the 3^rd^ exon of the PHD1 gene, leading to early termination of the PHD1 protein at amino acid 88. The *fwa* background RdDM mutants, including *nrpd1-4* (SALK_083051), *suvh2* (SALK_079574) *suvh9* (SALK_048033), *morc6-3* (GABI_599B06), *rdm1-4* (EMS)^54^, drd1-6 (EMS)^55^, *dms3-4* (SALK_125019C), *nrpe1-1* (EMS), and *drm1-2* (SALK_031705) *drm2-2* (SALK_150863) were previously described^43^. The other *fwa* background mutants in MOM1 complex were *phd1-2, aipp3-1* (GABI_058D11), *mom1-3* (SALK_141293), *mom2-1* (WiscDsLox364H07), and *pial1* (CS358389) *pial2* (SALK_043892), which were generated by crossing *fwa-4* to corresponding mutants. F2 offspring plants with late flowering phenotype were genotyped for homozygous T-DNA mutant alleles, and propagated to F3 generation. Then, F3 populations were screened for non-segregating homogenous late flowering phenotype. For IP-MS comparisons of MOM1-FLAG in *mom1-7* mutant background, to that in the backgrounds of *aipp3-1, mom2-2*, as well as *aipp3/mom2-2* double mutants, MOM1-FLAG transgenic lines were constructed by recombineering 2xYpet-3xFLAG encoding DNA sequence in frame with the C terminus of MOM1 gene, in a transformation-competent artificial chromosome clone (JAtY68M20 (68082 bp)) using a bacterial recombineering approach^56^ and transformed into *mom1-7* mutants. Then this MOM1-FLAG transgenic line was crossed into *aipp3-1, mom2-2*, as well as *aipp3/mom2-2* double mutant backgrounds. For transgenic plants of FLAG epitope tagged, MYC epitope tagged and ZF tagged proteins used in all other IP-MS, ChIP-seq and ZF tethering experiments, genomic DNA fragments including the promoter region were amplified and cloned into entry vectors (pENTR-D or PCR8 from Invitrogen) and cloned into destination vectors with C-terminal 3xFLAG (pEG302_GW_3xFLAG), MYC (pEG302_GW_9xMYC) and ZF108 (pEG302_GW_3xFLAG_ZF108) by LR clonase II (Invitrogen). Primers used in this study were listed in Supplementary Table 4. Agrobacterium mediated floral dipping (strain Agl0) were used to generate transgenic plants in corresponding loss-of-function mutant backgrounds or specific mutant backgrounds as indicated.

### IP-MS and cross-linking IP-MS

Native IP-MS and cross-linking IP-MS were performed following a method described in a recent paper with modifications^40^. 50 ml of liquid nitrogen flash-frozen unopened flower buds from FLAG epitope tagged transgenic plants were used for each IP-MS experiment and flower buds of Col-0 plants were used as control. Flower tissue were ground to fine powder in liquid nitrogen with Retsch homogenizer. For Native IP-MS, tissue powder was resuspended in 25 ml IP buffer (50 mM Tris-HCl pH 8.0, 150 mM NaCl, 5 mM EDTA, 10% glycerol, 0.1% Tergitol, 0.5 mM DTT, 1 mg/ml Pepstatin A, 1 mM PMSF, 50 uM MG132 and cOmplete EDTA-free Protease Inhibitor Cocktail (Roche)) and further homogenized with dounce homogenizer. The lysates were filtered with Miracloth and centrifuged at 20,000 g for 10 min at 4 °C. The supernatant was incubated with 250 μL anti-FLAG M2 magnetic beads (Sigma) at 4 °C for 2 hours with constant rotation. The magnetic beads were washed with IP buffer and eluted with 250 ug/ml 3xFLAG peptides. Eluted proteins were used for Trichloroacetic acid (TCA) precipitation and mass spectrometric analysis.

For Crosslinking IP-MS, flower tissue powder was resuspended in 40 ml nuclei extraction buffer^40^ with 1.5 mM EGS (Ethylene Glyco-bis (succinimidylsuccinate)) and rotated at room temperature for 10 min. Then the lysate was supplemented with formaldehyde at 1% final concentration and rotated at room temperature for another 10 min followed by adding glycine to stop crosslinking. The crosslinked lysate was filtered through Miracloth and centrifuged for 20 min at 2880 g. The pellet (which contains the nuclei) was resuspended in 3 ml of extraction buffer 2 (0.25 M sucrose, 10 mM Tris-HCl pH 8.0, 10 mM MgCl_2_, 1% Triton X-100, 5 mM 2-Mercaptoethanol, 0.1 mM PMSF, 5mM Benzamidine and cOmplete EDTA-free Protease Inhibitor Cocktail (Roche)), then centrifuged at 12,000 g for 10 min at 4 °C. Then, the pellet was carefully resuspended in 1.2ml nuclear lysis buffer (50 mM Tris-HCl pH 8.0, 10 mM EDTA, 1% SDS, 0.1 mM PMSF, 5 mM Benzamidine and cOmplete EDTA-free Protease Inhibitor Cocktail (Roche)) and incubated on ice for 10 min. After that, 5.1 ml dilution buffer (1.1% Triton x-100, 1.2 mM EDTA, 16.7 mM Tris-HCl pH 8.0, 167 mM NaCl, 1 mM PMSF, 5 mM Benzamidine and cOmplete EDTA-free Protease Inhibitor Cocktail (Roche)) was added and mixed by pipetting. Resuspended nuclei were split into 3x 2.1ml aliquots for sonication of 22 min (30 s on/30s off) with Bioruptor Plus (Diagenode). Sheared lysate from the same sample was combined and centrifuged at 12,000 g for 10 min at 4 °C. Another 6 ml of dilution buffer and 250 μL anti-FLAG M2 magnetic beads (Sigma) were added to the supernatant and the sample was incubated at 4 °C for 2 hours with constant rotation. Then, the magnetic beads were washed and eluted with 250 ug/ml 2xFLAG peptides. Eluted protein was used for Trichloroacetic acid (TCA) precipitation and mass spectrometric analysis.

The mass spectrometry procedure were performed as previously reported^40^. MS/MS database searching was performed using MaxQuant (1.6.10.43) against newest *Arabidopsis thaliana* proteome database from http://www.uniprot.org. Analysis of raw data was obtained from the LC–MS runs using MaxQuant with the integrated Andromeda peptide search engine using default setting with enabled LFQ normalization. Data sets were filtered at a 1% FDR at both the PSM and protein levels. The MaxQuant peptide intensity and MS/MS counts were used for all peptide quantitation. For Fig. 1d, fold of change of MS/MS counts and P value of MOM1-FLAG lines crosslinking IP-MS compared to crosslinking IP-MS of Col-0 control were calculated by LIMMA^57^.

### Chromatin immunoprecipitation sequencing (ChIP-seq)

We followed previous protocol^28,40^ for ChIP-seq with some modifications. Briefly, 15 ml of unopened flower buds were collected for each ChIP and flash-frozen in liquid nitrogen. The flower tissue was ground to fine powder with Retsch homogenizer in liquid nitrogen and resuspended in nuclei extraction buffer (50 mM HEPES pH 8.0, 1 M sucrose, 5 mM KCl, 5 mM MgCl_2_, 0.6% Triton X-100, 0.4 mM PMSF, 5 mM benzamidine, cOmplete EDTA-free Protease Inhibitor Cocktail (Roche), 50uM MG132). For transgenic lines of MOM1-MYC in *mom1-7* and PIAL2-MYC in *pial2-1*, EGS was first added to resuspended lysate to 1.5 mM and the tissue lysate was incubated at room temperature for 10 min with rotation. Then the lysate was supplemented with formaldehyde at 1% and rotated at room temperature for another 10 min followed by adding glycine to stop crosslinking. For ChIP of all other proteins, crosslinking was performed by directly supplementing formaldehyde to 1% without adding EGS, then rotated at room temperature for 10 min followed by adding glycine to stop crosslinking. The crosslinked nuclei were isolated, lysed with Nuclei Lysis Buffer and diluted with ChIP Dilution Buffer as previously described^40^. Then the lysate was sonicated for 22 min (30 s on/30s off) with Bioruptor Plus (Diagenode). After centrifugation, antibody for FLAG epitope (M2 monoclonal antibody, Sigma F1804, 10 ul per ChIP) or for MYC epitope (Cell Signaling, 71D10, 20 ul per ChIP) were added to the supernatant and incubated at 4 °C overnight with rotation. Then, Protein A and Protein G Dynabeads (Invitrogen) were added and incubated at 4 °C for 2 hours with rotation. After that, as previously described^40^, the beads were washed and eluted, and the eluted chromatin was reverse-crosslinked by adding 20 ul 5 M NaCl and incubated at 65 °C overnight followed by treatment of Proteinase K (Invitrogen) for 4 hours at 45 °C. DNA was purified and precipitated with 3 M Sodium Acetate, GlycoBlue (Invitrogen) and ethanol at -20 °C overnight. After centrifugation, the precipitated DNA was washed with ice cold 70% ethanol, air dried and dissolved in 120 ul of H_2_O. ChIP-seq libraries were prepared with Ovation Ultra Low System V2 kit (NuGEN), and sequenced on Illumina NovaSeq 6000 or HiSeq 4000 instruments.

For ChIP-seq analysis, raw reads were trimmed using trim_galore (https://www.bioinformatics.babraham.ac.uk/projects/trim_galore/) and aligned to the TAIR10 reference genome with bowtie2 (v2.4.2)^58^ allowing zero mismatch and reporting one valid alignment for each read. The Samtools (v1.15)^59^ were used to convert sam files to bam files, sort bam files and remove duplicate reads. Track files in bigWig format were generated using bamCoverage of deeptools (v3.5.1)^60^ with RPKM normalization. Peaks were called with MACS2 (v2.1.2)^61^ and peaks frequently identified in previous ChIP-seq of Col-0 plant with M2 antibody for FLAG epitope were removed from analysis.

For unsupervised clustering of Pol V and MOM1 peaks (Fig. 6b), RPKM of Pol V^34^, MOM1 and corresponding control ChIP-seqs over merged peaks of Pol V and MOM1 were calculated with custom scripts. Then, log2(PolV RPKM/control RPKM) and log2(MOM1 RPKM /control RPKM) were calculated and used for unsupervised clustering with the ConcensusClusterPlus R package (v1.60.0)^62^. For analysis of ChIP signal over TEs located in euchromatic arms versus TEs located in pericentromeric regions (Fig. 6a), the pericentromeric regions were as previously defined^63^.

### RNA sequencing

For RNA-seq experiments, twelve-day old seedlings grown on half MS medium (Murashige and Skoog Basal Medium) were collected and flash-frozen in liquid nitrogen. RNA was extracted with Direct-zol RNA MiniPrep kit (Zymo Research) and 1ug of total RNA was used to prepare RNA-seq libraries with TruSeq Stranded mRNA kit (Illumina), and the libraries were sequenced on Illumina NovaSeq 6000 instruments.

The raw reads of RNA-seq were aligned to the TAIR10 reference genome with bowtie2. Rsem-calculate-expression from RSEM^64^ with default settings was used to calculate expression levels. DEGs and DE-TEs were calculated with run_DE_analysis.pl from Trinity version 2.8.5^65^ and log2 FC ≥ 1 and FDR < 0.05 were used as the cut off. RNA-seq track files in bigWig format were generated using bamCoverage of deeptools (v3.1.3) with RPKM normalization.

### Whole Genome Bisulfite Sequencing

Rosette leaves of about one-month-old *Arabidopsis* Col-0 wild type, *phd1-2, phd1-3, mom2-2, aipp3-1, fwa* plants and ZF transgenic lines (MOM1-ZF, MOM2-ZF, PIAL1-ZF, PIAL2-ZF and PHD1-ZF) T2 plants with early flowering phenotype were collected for DNA extraction using DNeasy Plant Mini Kit (QIAGEN). 500 ng DNA was sheared with Covaris S2 (Covaris) into around 200bp at 4°C. The DNA fragments were used to perform end repair reaction using the Kapa Hyper Prep kit (Roche), and together with Illumina TruSeq DNA sgl Index Set A/B (Illumina) to perform adapter ligation. The ligation products were purified with AMPure beads (Beckman Coulter), and then converted with EpiTect Bisulfite kit (QIAGEN). The converted ligation products were used as templates, together with the primers from the Kapa Hyper Prep kit (Roche) and MyTaq Master mix (Bioline) to perform PCR. The PCR products were purified with AMPure beads (Beckman Coulter) and sequenced by Illumina NovaSeq 6000 instrument.

The WGBS data analysis has been was performed as previously described^43^ with minor modifications. The WGBS raw reads were aligned to both strands of the TAIR10 reference genome using BSMAP (v.2.74)^66^, allowing up to 2 mismatches and 1 best hit. Reads with more than 3 consecutives methylated CHH sites were removed, and the methylation level was calculated with the ratio of C/(C+T). For Fig. 2d, the methylation levels at 1kb flanking regions of ZF off target sites^43^ in MOM1-ZF, MOM2-ZF, PIAL1-ZF, PIAL2-ZF and PHD1-ZF were subtracted by the methylation level of *fwa* and plotted with R package pheatmap.

For Fig. 5a, the hcDMRs (p < 0.01, > 33 supported controls) of Col-0 wild type, *aipp3-1, phd1-2, mom1-2, mom2-1, pial1 pial2, morc6*, and *morchex* mutants were called using a previous method^8^, which were then used to generate the heat map using R package pheatmap [R. Kolde, Pheatmap: pretty heatmaps]. For Fig. 5b, Overlap Enrichment was calculated by using HOMER^67^ mergePeaks to identify overlapped CHH hcDMR regions and followed by normalization with genome size and over random shuffles.

### BS-PCR-seq

Rosette leaves of about one-month-old plants were collected and subject to DNA extraction with CTAB method followed by bisulfite DNA conversion using the EpiTect Bisulfite kit (QIAGEN) kit. Three regions of the *FWA* gene were amplified from the converted DNA with Pfu Turbo Cx (Agilent): Region 1 (chr4: 13038143-13038272), Region 2 (chr4: 13038356-13038499) and Region3 (chr4: 13038568-13038695). Primers used are listed in Supplementary Table 4. Libraries were prepared with the purified PCR product by the Kapa DNA Hyper Kit (Roche) together with TruSeq DNA UD indexes for Illumina (Illumina) and were sequenced on Illumina iSeq 100 or HiSeq 4000 instruments.

BS-PCR-seq data was analyzed as previously described^43^. Briefly, raw reads were aligned to both strands of the TAIR10 reference genome with BSMAP (v.2.90)^66^ allowing up to 2 mismatches and 1 best hit. After quality filtering, the methylation level of cytosines was calculated as the ratio of C/(C+T), and customized R scripts were used to plot methylation data over the *FWA* region 1-3.

### ATAC-seq

Fresh unopened flower buds of about one-month-old Col-0 and *mom1-3* mutant plants were collected for nuclei extraction and ATAC-seq, with two replicates for each genotype. The nuclei collection process from unopened flower buds is as described previously^34^. Freshly isolated nuclei were used for ATAC-seq as described elsewhere^68^. Unopened flower buds were collected for extraction of nuclei as follows. About 5 grams of unopened flower buds was collected and immediately transferred into ice-cold grinding buffer (300 mM sucrose, 20 mM Tris pH 8, 5 mM MgCl_2_, 5 mM KCl, 0.2% Triton X-100, 5 mM β-mercaptoethanol, and 35% glycerol). The samples were ground with Omni International General Laboratory Homogenizer on ice and then filtered through a two-layer Miracloth and a 40-µm nylon mesh Cell Strainer (Fisher). Samples were spin filtered for 10 min at 3,000 *g*, the supernatant was discarded, and the pellet was resuspended with 25 ml of grinding buffer using a Dounce homogenizer. The wash step was performed twice in total, and nuclei were resuspended in 0.5 ml of freezing buffer (50 mM Tris pH 8, 5 mM MgCl_2_, 20% glycerol, and 5 mM β-mercaptoethanol). Nuclei were subjected to a transposition reaction with Tn5 (Illumina). For the transposition reaction, 25 µl of 2x DMF (66 mM Tris-acetate pH 7.8, 132 mM K-Acetate, 20 mM Mg-Acetate, and 32% DMF) was mixed with 2.5 µl Tn5 and 22.5 µl nuclei suspension at 37°C for 30 min. Transposed DNA fragments were purified with ChIP DNA Clean & Concentrator Kit (Zymo). Libraries were prepared with Phusion High-Fidelity DNA Polymerase (NEB) in a system containing 12.5 µl 2x Phusion, 1.25 µl 10 mM Ad1 primer, 1.25 µl 10 mM Ad2 primer, 4 µl ddH2O, and 6 µl purified transposed DNA fragments. The ATAC-seq libraries were sequenced on HiSeq 4000 platform (Illumina).

ATAC-seq data analysis was also performed as previously described^69^. Briefly, raw reads were adaptor-trimmed with trim_galore and mapped to the TAIR10 reference genome with Bowtie2^58^ (-X 2000 -m 1). After removing duplicate reads and reads mapped to chloroplast and mitochondrial, ATAC-Seq open chromatin peaks of each replicate were called using MACS2 with parameters -p 0.01 --nomodel --shift -100 --extsize 200. Consensus peaks between replicates were identified with bedtools (version 2.26.0) intersect and differential accessible peaks were called with the R packge edgeR^70^ (version 3.30.0). Merged bigwig file of the two replicates were used for heatmap and metaplot.

### RT-qPCR

Rossette leaves of about one-month-old plants were collected for RNA extraction with Zymo Direct-Zol RNA miniprep Kit (Zymo Research). 1 ug of RNA were used for cDNA synthesis with iScript cDNA Synthesis Kit (Bio-Rad). qPCR was performed with iQ SYBR Green Supermix (Bio-Rad) and primers for qPCR were listed in Supplementary Table 4.

### McrBC assay

Genomic DNA extracted with the CTAB method were treated with RNase A (Qiagen) and diluted to about 100 ng/ul. 10 ul of diluted DNA were used for McrBC digestion (NEB, 4 h at 37 °C) or mock digestion (the same volume of H_2_O instead of McrBC enzyme was added with all other components the same in the reaction, was also kept for 4 h at 37 °C). Relative undigested *FWA* promoter quantity (McrBC treated / H_2_O treated) was determined with qPCR and primers used were listed in Supplementary Table 4.

### Flowering time measurement

Total true leaf numbers (sum of rosette leaf number and cauline leaf number) after bolting of the plants were used as measurement of flowering time. Plants with less than 20 true leaf number were considered as early flowering. The numbers of independent plants (n) scored for each population are listed in Supplementary Table 5.

### Yeast two-hybrid (Y2H)

The cDNA sequences of PIAL1, PIAL2, MOM2, MORC6, and MOM1 CMM2 domain (aa1660-aa1860)^5^ were first cloned into gateway entry vectors followed by LR reaction with pGBKT7-GW (Addgene 61703) and pGADT7-GW (Addgene 61702) destination vectors. Pairs of plasmid DNA for the desired protein interaction to be tested were co-transformed into the yeast strain AH109. Combinations of the empty pGBKT7-GW or pGADT7-GW vectors and the plasmids of desired proteins were used for transformation of yeast cells to test for self-activation. Transformed yeast cells were plated on synthetic dropout medium without Trp and Leu (SD-TL) and incubated for 2-3 days to allow for the growth of positive colonies carrying both plasmids. Three yeast colonies of each tested protein interaction pairs were picked and mixed in 150μl 1xTE solution, and 3μl of the 1xTE solution with the yeast cells were blotted on synthetic dropout medium without Trp, Leu, and His (SD-TLH) and with 5mM 3-amino-1,2,4-triazole (3AT) to inhibit background growth. Growth of yeast on SD-TLH with 5mM 3AT medium after 2-3 days of incubation indicates the interaction between the GAL4-AD fusion protein and the GAL4-BD fusion protein.

### Co-immunoprecipitation

The Co-immunoprecipitation experiment was performed following previous protocol with some modifications^71^. 2 grams of 2-week-old seedling tissue were collected from MORC6-FLAG X PIAL2-Myc F1 generation and PIAL2-Myc transgenic plants and ground into fine powder in liquid nitrogen. The tissue powder was resuspended with 10 ml IP buffer, and incubated for 20 min at 4°C. Then the lysate was centrifuged and filtered with Miracloth twice. 30 μL of anti-FLAG M2 Affinity Gel (Millipore) was added to the supernatant and incubated for 2 hours at 4°C. Then, the anti-FLAG beads were washed with IP buffer for 5 times, and eluted with 40ul elution buffer (IP buffer with 100 ug/ml 3xFLAG peptide). The eluted protein was used for western blot.

## Data availability

All high-throughput sequencing data generated in this study are accessible at the National Center for Biotechnology information Gene Expression Omnibus via series accession GSE221679. (also weblink here https://www.ncbi.nlm.nih.gov/geo/query/acc.cgi?acc=GSE221679).

## Code availability

The customized code used in this manuscript can be distributed upon request. Requests should be addressed to S.E.J.

## Acknowledgements

We thank Suhua Feng and Mahnaz Akhavan for support with high-throughput sequencing at the UCLA Broad Stem Cell Research Center BioSequencing Core Facility. This work was supported by NIH R35 GM130272 to S.E.J. S.E.J is an Investigator of the Howard Hughes Medical Institute.

## Author contributions

Z.L, M.W., Z.Z and S.E.J. designed the research, interpreted the data, and wrote the manuscript; Z.L, M.W. and Z.Z performed experiments and performed bioinformatic data analysis; Y.J.A., and J.W performed IP-MS and interpreted the data. S.B. and J.A.L. contributed to gathering mutant materials, construction of transgenic lines, performing initial ZF108 tethering assays and discussions. J.G.B contributed to PHD1-ZF108 materials. S.F. performed BS-PCR-seq and high throughput sequencing; X.W provided technical support.

## Figure Legends

**Supplementary Fig. 1** | **Example of AIPP3 Group1 and Group2 ChIP-seq Peaks**. Screenshots of MOM1-Myc, PHD1-FLAG, AIPP3-FLAG and PHD3-FLAG ChIP-seq signals over representative AIPP3 Group1 peaks (**a**) and Group2 peaks (**b**), with control ChIP-seq signals subtracted.

**Supplementary Fig. 2** | **PIAL1-ZF and AIPP3-ZF silence *FWA* less efficiently than ZF tethering of other MOM1 complex components. a**, Flowering time of *fwa*, Col-0, and T1 populations of MOM1-ZF, MOM2-ZF, PIAL1-ZF, PIAL2-ZF and PHD1-ZF in the *fwa* background. **b**, left panel: qRT-PCR showing the relative mRNA level of *FWA* gene in PIAL1-ZF T1 plants in *fwa* background. Right panel: qPCR showing the relative *FWA* promoter DNA quantity after McrBC treatment in PIAL1-ZF T1 plants in *fwa* background. Bar plots and error bars indicate the mean and standard error of three technical replicates, respectively, with individual technical replicates shown as dots. **c**, Flowering time of *fwa*, Col-0, and representative T2 populations of AIPP3-ZF in *fwa* background. For **a** and **c**, the numbers of independent plants (n) scored for each population are listed in Supplementary Table 5. **d**, CG, CHG, and CHH DNA methylation levels over *FWA* promoter regions measured by BS-PCR-seq in Col-0, *fwa* and representative T2 plants of AIPP3-ZF with (+) or without (-) transgenes in the *fwa* background. Pink vertical boxes indicate ZF binding sites.

**Supplementary Fig. 3** | **ZF tethering of MOM1 complex components and miniMOM1 lead to DNA methylation. a**, Screenshots of Whole Genome Bisulfite Sequencing (WGBS) showing CG, CHG, and CHH DNA methylation level over a representative ZF off-target site in *fwa*, and representative T2 plants of MOM1-ZF, MOM2-ZF, PIAL1-ZF, PIAL2-ZF and PHD1-ZF in the *fwa* background. **b**, Flowering time of miniMOM1-ZF T1 plants in the *fwa* background (upper panel) and representative T2 lines (lower panel). The numbers of independent plants (n) scored for each population are listed in Supplementary Table 5. **c**, CG, CHG, and CHH DNA methylation levels over *FWA* promoter regions measured by BS-PCR-seq in Col-0, *fwa*, and representative mini-MOM1-ZF T2 plants with (+) or without (-) miniMOM1-ZF transgenes in the *fwa* background. Pink vertical boxes indicated ZF binding sites.

**Supplementary Fig. 4** | **Analysis of ZF tethering of MOM1 complex components and MORC6 in mutant backgrounds. a**, Flowering time of MOM1-ZF T1 plants in the backgrounds of *fwa* introgressed into *aipp3-1, phd1-2, mom2-1* and *pial1/2* mutants; Flowering time of PHD1-ZF T1 plants in the backgrounds of *fwa* introgressed into *aipp3-1, mom1-3* and *pial1/2* mutants. **b**, Flowering time of PIAL2-ZF T1 plants in the backgrounds of *fwa* introgressed into *aipp3-1, phd1-2, mom1-3* and *mom2-1*; Flowering time of MOM2-ZF T1 plants in the backgrounds of *fwa* introgressed into *aipp3-1, phd1-2, mom1-3* and *pial1/2*. **c**, Flowering time of *fwa* introgressed into *mom1-3, pial1/2* and *aipp3* plants, with Col-0 and *fwa* plants as controls. **d**, Flowering time of MOM2-ZF and PIAL1-ZF T1 plants in the background of *fwa* introgressed into *mom6-3*; Flowering time of MORC6-ZF T1 plants in the backgrounds of *fwa* introgressed into *mom1-3, mom2-1, pial1/2, phd1-2* and *aipp3-1*. The numbers of independent plants (n) scored for each population are listed in Supplementary Table 5.

**Supplementary Fig. 5** | **Flowering time of *FWA* transgene T1 plants in MOM1 complex component mutant backgrounds. a**, Comparison of the flowering time of T1 plant populations with *FWA* transgenes in the Col-0, *nrpe1-11*, MOM1 complex component mutant backgrounds. One-way ANOVA followed by Dunnett’s multiple comparison tests were used for statistical analysis. **b**, Flowering time of Col-0, *nrpe1-11, mom1-3, pial1/2, mom2-2, aipp3-1* and *phd1-2* plants. The numbers of independent plants (n) scored for each population are listed in Supplementary Table 5.

**Supplementary Fig. 6** | **RNA-seq analysis of the mutants of MOM1 complex components. a**, Dotplots showing the differentially expressed TEs (compared to Col-0 control) over the five *Arabidopsis* chromosomes in the *nrpe1-11, mom1-2, mom1-3, pial1/2, morc6-3, morchex, aipp3-2, aipp3-1, pial1-2, pial2-1, mom2-1, mom2-2, phd1-2* and *phd1-4* mutant backgrounds. Red and blue dots indicate upregulated and down regulated TEs in mutants compared to Col-0 control, respectively. The positions of pericentromeric heterochromatin regions of each chromosome are annotated at the bottom of each plot. **b**, Heatmap showing the expression level of differentially expressed TEs (DE TEs, n=423) in three replicates of *mom1-2, mom1-3, pial1/2, morc6-3, morchex, aipp3-1, aipp3-2, pial1-2, pial2-1, mom2-1, mom2-2, phd1-2* and *phd1-4* mutant plants versus Col-0 plants. Expression level of these TEs in *nrpe1-11* mutant and corresponding Col-0 control plants are also plotted for comparison.

## References

1. Law, J. A. & Jacobsen, S. E. Establishing, maintaining and modifying DNA methylation patterns in plants and animals. Nat. Rev. Genet. 11, 204–220 (2010).

2. Slotkin, R. K. & Martienssen, R. Transposable elements and the epigenetic regulation of the genome. Nat. Rev. Genet. 8, 272–285 (2007).

3. Amedeo, P., Habu, Y., Afsar, K., Mittelsten Scheid, O. & Paszkowski, J. Disruption of the plant gene MOM releases transcriptional silencing of methylated genes. Nature 405, 203–206 (2000).

4. Numa, H. et al. Transduction of RNA-directed DNA methylation signals to repressive histone marks in Arabidopsis thaliana. EMBO J. 29, 352–362 (2010).

5. Han, Y.-F. et al. The SUMO E3 Ligase-Like Proteins PIAL1 and PIAL2 Interact with MOM1 and Form a Novel Complex Required for Transcriptional Silencing. Plant Cell 28, 1215–1229 (2016).

6. Yokthongwattana, C. et al. MOM1 and Pol-IV/V interactions regulate the intensity and specificity of transcriptional gene silencing. EMBO J. 29, 340–351 (2010).

7. Moissiard, G. et al. Transcriptional gene silencing by Arabidopsis microrchidia homologues involves the formation of heteromers. Proc. Natl. Acad. Sci. U. S. A. 111, 7474–7479 (2014).

8. Zhang, Y. et al. Large-scale comparative epigenomics reveals hierarchical regulation of non-CG methylation in Arabidopsis. Proc. Natl. Acad. Sci. U. S. A. 115, E1069–E1074 (2018).

9. Feng, S. et al. Genome-wide Hi-C analyses in wild-type and mutants reveal high-resolution chromatin interactions in Arabidopsis. Mol. Cell 55, 694–707 (2014).

10. Mittelsten Scheid, O., Probst, A. V, Afsar, K. & Paszkowski, J. Two regulatory levels of transcriptional gene silencing in Arabidopsis. Proc. Natl. Acad. Sci. U. S. A. 99, 13659–13662 (2002).

11. Probst, A. V, Fransz, P. F., Paszkowski, J. & Mittelsten Scheid, O. Two means of transcriptional reactivation within heterochromatin. Plant J. 33, 743–749 (2003).

12. Caikovski, M. et al. Divergent evolution of CHD3 proteins resulted in MOM1 refining epigenetic control in vascular plants. PLoS Genet. 4, e1000165 (2008).

13. Nishimura, T. et al. Structural basis of transcriptional gene silencing mediated by Arabidopsis MOM1. PLoS Genet. 8, e1002484 (2012).

14. Zhao, Q.-Y. & He, X.-J. Exploring potential roles for the interaction of MOM1 with SUMO and the SUMO E3 ligase-like protein PIAL2 in transcriptional silencing. PLoS One 13, e0202137 (2018).

15. Chan, S. W.-L. et al. RNA silencing genes control de novo DNA methylation. Science 303, 1336 (2004).

16. Matzke, M. A. & Mosher, R. A. RNA-directed DNA methylation: an epigenetic pathway of increasing complexity. Nat. Rev. Genet. 15, 394–408 (2014).

17. Matzke, M. A., Kanno, T. & Matzke, A. J. M. RNA-Directed DNA Methylation: The Evolution of a Complex Epigenetic Pathway in Flowering Plants. Annu. Rev. Plant Biol. 66, 243–267 (2015).

18. Wendte, J. M. & Pikaard, C. S. The RNAs of RNA-directed DNA methylation. Biochim. Biophys. acta. Gene Regul. Mech. 1860, 140–148 (2017).

19. Law, J. A., Vashisht, A. A., Wohlschlegel, J. A. & Jacobsen, S. E. SHH1, a homeodomain protein required for DNA methylation, as well as RDR2, RDM4, and chromatin remodeling factors, associate with RNA polymerase IV. PLoS Genet. 7, e1002195 (2011).

20. Zhou, M., Palanca, A. M. S. & Law, J. A. Locus-specific control of the de novo DNA methylation pathway in Arabidopsis by the CLASSY family. Nat. Genet. 50, 865–873 (2018).

21. Blevins, T. et al. Identification of Pol IV and RDR2-dependent precursors of 24 nt siRNAs guiding de novo DNA methylation in Arabidopsis. Elife 4, e09591 (2015).

22. Zhai, J. et al. A One Precursor One siRNA Model for Pol IV-Dependent siRNA Biogenesis. Cell 163, 445–455 (2015).

23. Henderson, I. R. et al. Dissecting Arabidopsis thaliana DICER function in small RNA processing, gene silencing and DNA methylation patterning. Nat. Genet. 38, 721–725 (2006).

24. Stroud, H., Greenberg, M. V. C., Feng, S., Bernatavichute, Y. V. & Jacobsen, S. E. Comprehensive analysis of silencing mutants reveals complex regulation of the Arabidopsis methylome. Cell 152, 352–364 (2013).

25. Lahmy, S. et al. Evidence for ARGONAUTE4-DNA interactions in RNA-directed DNA methylation in plants. Genes Dev. 30, 2565–2570 (2016).

26. McCue, A. D. et al. ARGONAUTE 6 bridges transposable element mRNA-derived siRNAs to the establishment of DNA methylation. EMBO J. 34, 20–35 (2015).

27. Olmedo-Monfil, V. et al. Control of female gamete formation by a small RNA pathway in Arabidopsis. Nature 464, 628–632 (2010).

28. Johnson, L. M. et al. SRA-and SET-domain-containing proteins link RNA polymerase v occupancy to DNA methylation. Nature 507, 124–128 (2014).

29. Johnson, L. M., Law, J. A., Khattar, A., Henderson, I. R. & Jacobsen, S. E. SRA-domain proteins required for DRM2-mediated de novo DNA methylation. PLoS Genet. 4, e1000280 (2008).

30. Liu, Z.-W. et al. The SET domain proteins SUVH2 and SUVH9 are required for Pol V occupancy at RNA-directed DNA methylation loci. PLoS Genet. 10, e1003948 (2014).

31. Wongpalee, S. P. et al. CryoEM structures of Arabidopsis DDR complexes involved in RNA-directed DNA methylation. Nat. Commun. 10, 3916 (2019).

32. Wierzbicki, A. T., Haag, J. R. & Pikaard, C. S. Noncoding Transcription by RNA Polymerase Pol IVb/Pol V Mediates Transcriptional Silencing of Overlapping and Adjacent Genes. Cell 135, 635–648 (2008).

33. Wierzbicki, A. T., Ream, T. S., Haag, J. R. & Pikaard, C. S. RNA polymerase V transcription guides ARGONAUTE4 to chromatin. Nat. Genet. 41, 630–634 (2009).

34. Liu, W. et al. RNA-directed DNA methylation involves co-transcriptional small-RNA-guided slicing of polymerase V transcripts in Arabidopsis. Nat. plants 4, 181–188 (2018).

35. Zhong, X. et al. Molecular mechanism of action of plant DRM de novo DNA methyltransferases. Cell 157, 1050–1060 (2014).

36. Moissiard, G. et al. MORC family ATPases required for heterochromatin condensation and gene silencing. Science 336, 1448–1451 (2012).

37. Jing, Y. et al. SUVH2 and SUVH9 Couple Two Essential Steps for Transcriptional Gene Silencing in Arabidopsis. Mol. Plant 9, 1156–1167 (2016).

38. Liu, Z.-W. et al. Two Components of the RNA-Directed DNA Methylation Pathway Associate with MORC6 and Silence Loci Targeted by MORC6 in Arabidopsis. PLoS Genet. 12, e1006026 (2016).

39. Harris, C. J. et al. Arabidopsis AtMORC4 and AtMORC7 Form Nuclear Bodies and Repress a Large Number of Protein-Coding Genes. PLoS Genet. 12, e1005998 (2016).

40. Xue, Y. et al. Arabidopsis MORC proteins function in the efficient establishment of RNA directed DNA methylation. Nat. Commun. 12, 4292 (2021).

41. Kinoshita, T. et al. One-way control of FWA imprinting in Arabidopsis endosperm by DNA methylation. Science 303, 521–523 (2004).

42. Soppe, W. J. J. et al. The late flowering phenotype of fwa mutants is caused by gain-of-function epigenetic alleles of a homeodomain gene. Mol. Cell 6, 791–802 (2000).

43. Gallego-Bartolomé, J. et al. Co-targeting RNA Polymerases IV and V Promotes Efficient De Novo DNA Methylation in Arabidopsis. Cell 176, 1068-1082.e19 (2019).

44. Qian, F. et al. A histone H3K27me3 reader cooperates with a family of PHD finger-containing proteins to regulate flowering time in Arabidopsis. J. Integr. Plant Biol. 63, 787–802 (2021).

45. Duan, C.-G. et al. A protein complex regulates RNA processing of intronic heterochromatin-containing genes in Arabidopsis. Proc. Natl. Acad. Sci. U. S. A. 114, E7377–E7384 (2017).

46. Zhang, Y.-Z. et al. Coupling of H3K27me3 recognition with transcriptional repression through the BAH-PHD-CPL2 complex in Arabidopsis. Nat. Commun. 11, 6212 (2020).

47. Moissiard, G. et al. Transcriptional gene silencing by Arabidopsis microrchidia homologues involves the formation of heteromers. Proc. Natl. Acad. Sci. U. S. A. 111, 7474–7479 (2014).

48. Greenberg, M. V. C. et al. Identification of genes required for de novo DNA methylation in Arabidopsis. Epigenetics 6, 344–354 (2011).

49. Habu, Y. et al. Epigenetic regulation of transcription in intermediate heterochromatin. EMBO Rep. 7, 1279–1284 (2006).

50. Huettel, B. et al. Endogenous targets of RNA-directed DNA methylation and Pol IV in Arabidopsis. EMBO J. 25, 2828–2836 (2006).

51. Liu, Q. et al. The characterization of Mediator 12 and 13 as conditional positive gene regulators in Arabidopsis. Nat. Commun. 11, 2798 (2020).

52. Zhong, Z. et al. MORC proteins regulate transcription factor binding by mediating chromatin compaction in active chromatin regions. bioRxiv 2022.11.01.514783 (2022) doi:10.1101/2022.11.01.514783.

53. Yan, L. et al. High-Efficiency Genome Editing in Arabidopsis Using YAO Promoter-Driven CRISPR/Cas9 System. Mol. Plant 8, 1820–1823 (2015).

54. Gao, Z. et al. An RNA polymerase II- and AGO4-associated protein acts in RNA-directed DNA methylation. Nature 465, 106–109 (2010).

55. Kanno, T. et al. Involvement of putative SNF2 chromatin remodeling protein DRD1 in RNA-directed DNA methylation. Curr. Biol. 14, 801–805 (2004).

56. Zhou, R., Benavente, L. M., Stepanova, A. N. & Alonso, J. M. A recombineering-based gene tagging system for Arabidopsis. Plant J. 66, 712–723 (2011).

57. Ritchie, M. E. et al. limma powers differential expression analyses for RNA-sequencing and microarray studies. Nucleic Acids Res. 43, e47 (2015).

58. Langmead, B. & Salzberg, S. L. Fast gapped-read alignment with Bowtie 2. Nat. Methods 9, 357–359 (2012).

59. Li, H. et al. The Sequence Alignment/Map format and SAMtools. Bioinformatics 25, 2078–2079 (2009).

60. Ramírez, F. et al. deepTools2: a next generation web server for deep-sequencing data analysis. Nucleic Acids Res. 44, W160–5 (2016).

61. Zhang, Y. et al. Model-based analysis of ChIP-Seq (MACS). Genome Biol. 9, R137 (2008).

62. Wilkerson, M. D. & Hayes, D. N. ConsensusClusterPlus: a class discovery tool with confidence assessments and item tracking. Bioinformatics 26, 1572–1573 (2010).

63. Bourguet, P. et al. The histone variant H2A.W and linker histone H1 co-regulate heterochromatin accessibility and DNA methylation. Nat. Commun. 12, 2683 (2021).

64. Li, B. & Dewey, C. N. RSEM: accurate transcript quantification from RNA-Seq data with or without a reference genome. BMC Bioinformatics 12, 323 (2011).

65. Grabherr, M. G. et al. Full-length transcriptome assembly from RNA-Seq data without a reference genome. Nat. Biotechnol. 29, 644–652 (2011).

66. Xi, Y. & Li, W. BSMAP: whole genome bisulfite sequence MAPping program. BMC Bioinformatics 10, 232 (2009).

67. Heinz, S. et al. Simple combinations of lineage-determining transcription factors prime cis-regulatory elements required for macrophage and B cell identities. Mol. Cell 38, 576–589 (2010).

68. Buenrostro, J. D., Giresi, P. G., Zaba, L. C., Chang, H. Y. & Greenleaf, W. J. Transposition of native chromatin for fast and sensitive epigenomic profiling of open chromatin, DNA-binding proteins and nucleosome position. Nat. Methods 10, 1213–1218 (2013).

69. Zhong, Z. et al. DNA methylation-linked chromatin accessibility affects genomic architecture in Arabidopsis. Proc. Natl. Acad. Sci. U. S. A. 118, (2021).

70. Robinson, M. D., McCarthy, D. J. & Smyth, G. K. edgeR: a Bioconductor package for differential expression analysis of digital gene expression data. Bioinformatics 26, 139–140 (2010).

71. Wang, Q. et al. Photoactivation and inactivation of Arabidopsis cryptochrome 2. Science 354, 343–347 (2016).

