## Supplementary figures for "The MOM1 complex recruits the RdDM machinery via MORC6 to establish *de novo* DNA methylation"

[2] Present address: Instituto de Biología Molecular y Celular de Plantas (IBMCP), CSIC-Universitat Politècnica de València, 46022 Valencia, Spain.

[5] Present address: Department of Plant and Microbial Biology, University of Zurich, CH-8008 Zurich Switzerland.

Supplementary Fig. 1- Supplementary Fig. 6

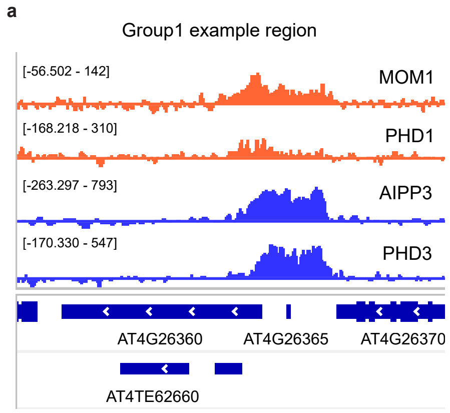


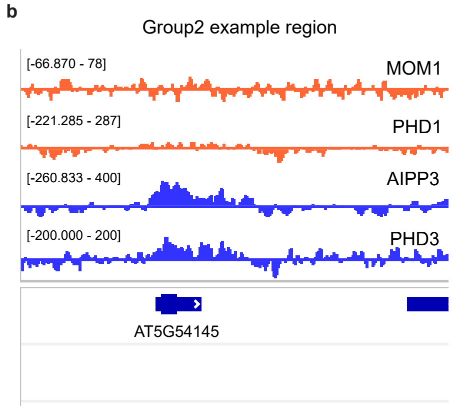


**Supplementary Fig. 1 | Example of AIPP3 Group1 and Group2 ChIP-seq Peaks.** Screenshots of MOM1-Myc, PHD1-FLAG, AIPP3-FLAG and PHD3-FLAG ChIP-seq signals over representative AIPP3 Group1 peaks (**a**) and Group2 peaks (**b**), with control ChIP-seq signals subtracted.

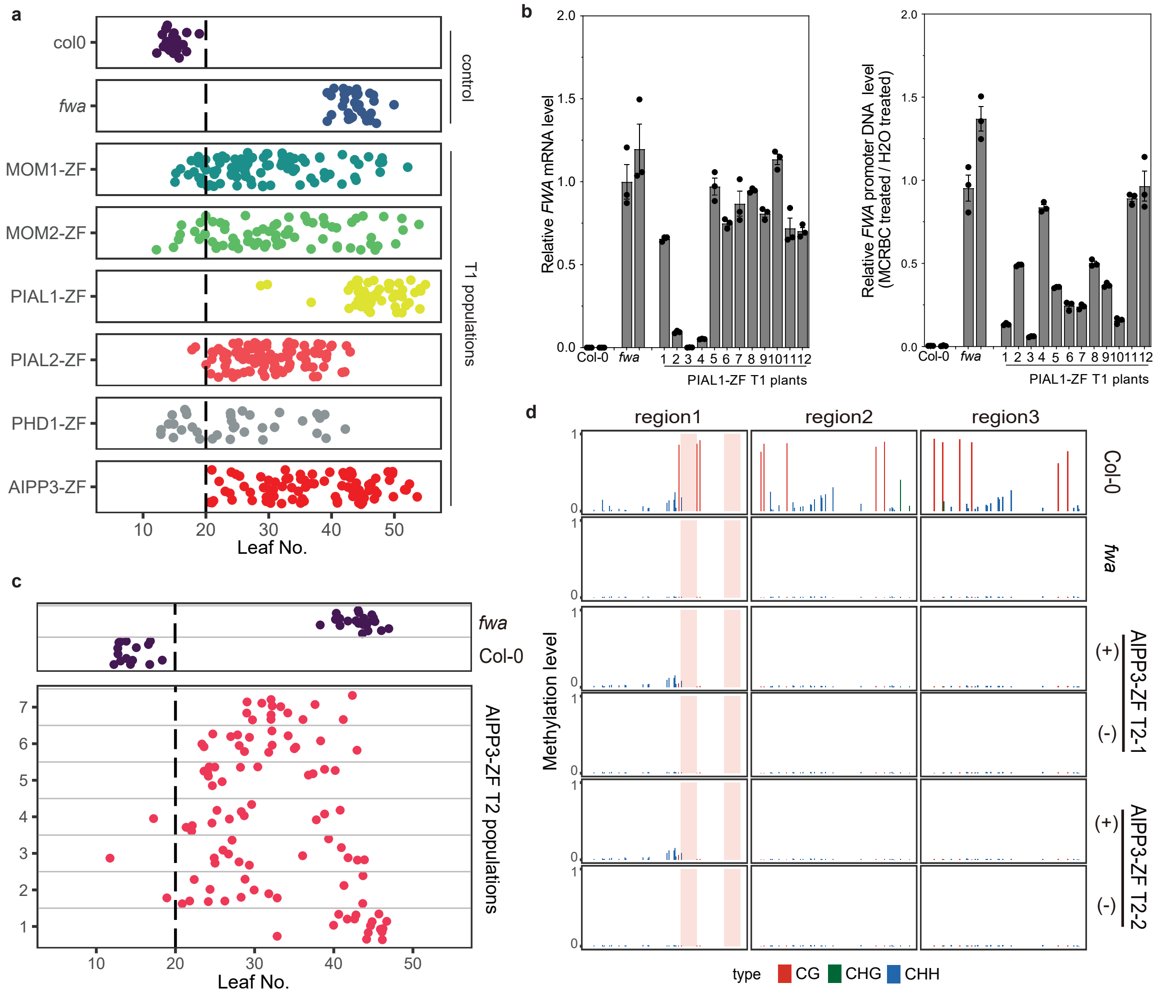


**Supplementary Fig. 2 | PIAL1-ZF and AIPP3-ZF silence *FWA* less efficiently than ZF tethering of other MOM1 complex components. a,** Flowering time of *fwa*, Col-0, and T1 populations of MOM1-ZF, MOM2-ZF, PIAL1-ZF, PIAL2-ZF and PHD1-ZF in the *fwa* background. **b**, left panel: qRT-PCR showing the relative mRNA level of *FWA* gene in PIAL1-ZF T1 plants in fwa background. Right panel: qPCR showing the relative FWA promoter DNA quantity after McrBC treatment in PIAL1-ZF T1 plants in fwa background. Bar plots and error bars indicate the mean and standard error of three technical replicates, respectively, with individual technical replicates shown as dots. **c**, Flowering time of *fwa*, Col-0, and representative T2 populations of AIPP3-ZF in fwa background. For **a** and **c**, the numbers of independent plants (n) scored for each population are listed in Supplementary Table 5. **d**, CG, CHG, and CHH DNA methylation levels over FWA promoter regions measured by BS-PCR-seq in Col-0, *fwa* and representative T2 plants of AIPP3-ZF with (+) or without (-) transgenes in the fwa background. Pink vertical boxes indicate ZF binding sites.

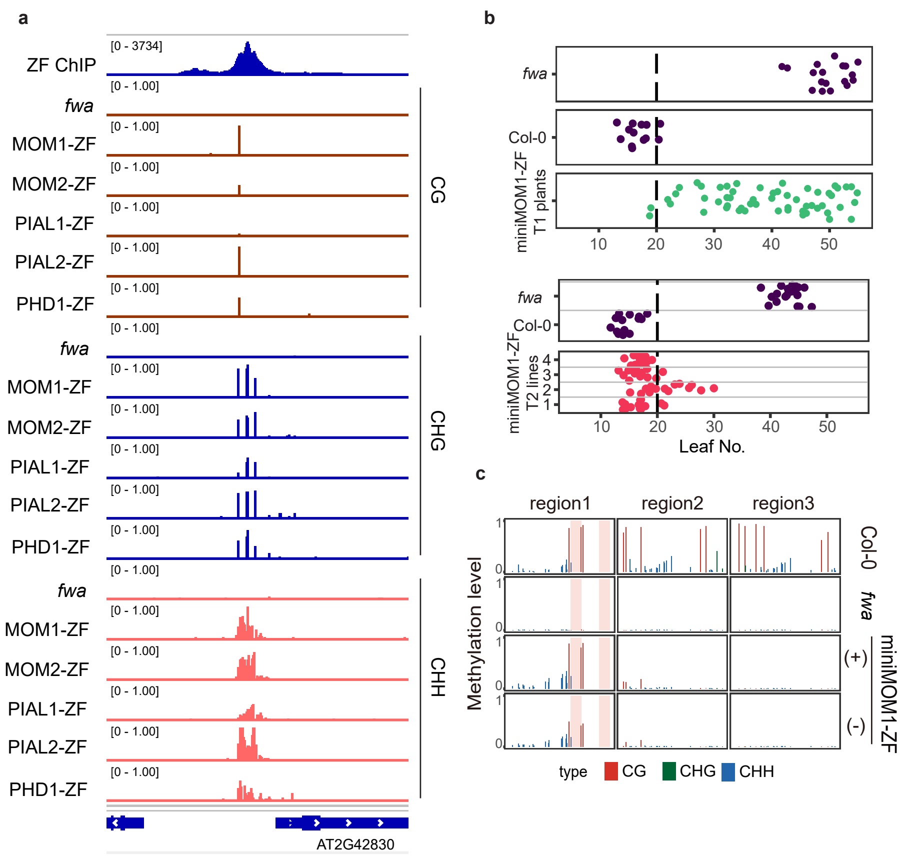


**Supplementary Fig. 3 | ZF tethering of MOM1 complex components and miniMOM1 lead to DNA methylation. a,** Screenshots of Whole Genome Bisulfite Sequencing (WGBS) showing CG, CHG, and CHH DNA methylation level over a representative ZF off-target site in *fwa*, and representative T2 plants of MOM1-ZF, MOM2-ZF, PIAL1-ZF, PIAL2-ZF and PHD1-ZF in the *fwa* background. **b**, Flowering time of miniMOM1-ZF T1 plants in the *fwa* background (upper panel) and representative T2 lines (lower panel). The numbers of independent plants (n) scored for each population are listed in Supplementary Table 5. **c**, CG, CHG, and CHH DNA methylation levels over FWA promoter regions measured by BS-PCR-seq in Col-0, *fwa*, and representative mini-MOM1-ZF T2 plants with (+) or without (-) miniMOM1-ZF transgenes in the *fwa* background. Pink vertical boxes indicated ZF binding sites.

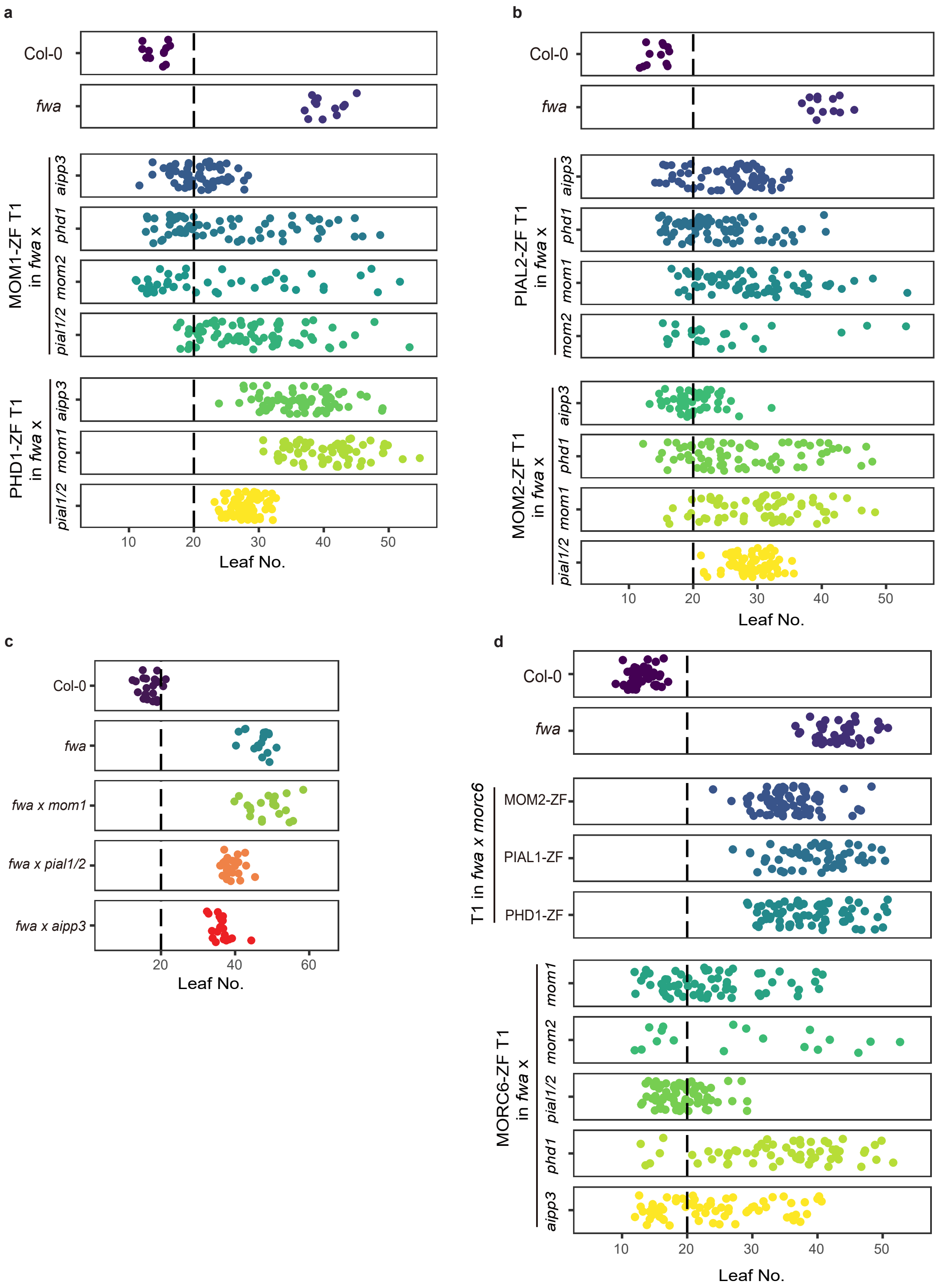


**Supplementary Fig. 4 | Analysis of ZF tethering of MOM1 complex components and MORC6 in mutant backgrounds. a,** Flowering time of MOM1-ZF T1 plants in the backgrounds of *fwa* introgressed into *aipp3-1, phd1-2, mom2-1* and *pial1/2* mutants; Flowering time of PHD1-ZF T1 plants in the backgrounds of *fwa* introgressed into *aipp3-1, mom1-3* and *pial1/2* mutants. **b**, Flowering time of PIAL2-ZF T1 plants in the backgrounds of *fwa* introgressed into *aipp3-1, phd1-2, mom1-3* and *mom2-1*; Flowering time of MOM2-ZF T1 plants in the backgrounds of *fwa* introgressed into *aipp3-1, phd1-2, mom1-3* and *pial1/2*. **c**, Flowering time of *fwa* introgressed into *mom1-3, pial1/2* and *aipp3* plants, with Col-0 and *fwa* plants as controls. **d**, Flowering time of MOM2-ZF and PIAL1-ZF T1 plants in the background of *fwa* introgressed into *mom6-3*; Flowering time of MORC6-ZF T1 plants in the backgrounds of *fwa* introgressed into *mom1-3, mom2-1, pial1/2, phd1-2* and *aipp3-1*. The numbers of independent plants (n) scored for each population are listed in Supplementary Table 5.

**a**

Comparison of flowering time as counted by leaf number of *FWA* transgene T1 plant populations in the following background:

| background | Leaf No. Mean¹ | n | Mean Difference² | Adjusted P Value | P value summary |
|---|---|---|---|---|---|
| Col-0 | 15.91 | 127 | 0 | NA | NA |
| *nrpe1-11* | 33.81 | 16 | -17.9 | <0.0001 | **** |
| *mom1-3* | 27.55 | 31 | -11.64 | <0.0001 | **** |
| *pial1/2* | 31.98 | 59 | -16.07 | <0.0001 | **** |
| *mom2-2* | 18.36 | 80 | -2.449 | 0.1055 | ns |
| *aipp3-1* | 13.25 | 104 | 2.663 | 0.0358 | * |
| *phd1-2* | 16.85 | 73 | -0.9359 | 0.9289 | ns |

¹ Leaf No. Mean = mean of (rosette + cauline leaf number)

² Mean difference = (mean leaf number in Col-0) - (mean leaf number in other background)

**b**

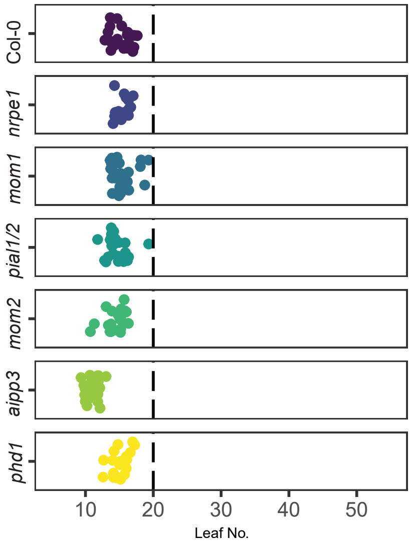


**Supplementary Fig. 5 | Flowering time of *FWA* transgene T1 plants in MOM1 complex component mutant backgrounds. a,** Comparison of the flowering time of T1 plant populations with *FWA* transgenes in the Col-0, *nrpe1-11*, MOM1 complex component mutant backgrounds. One-way ANOVA followed by Dunnett's multiple comparison tests were used for statistical analysis. **b**, Flowering time of Col-0, *nrpe1-11, mom1-3, pial1/2, mom2-2, aipp3-1* and *phd1-2* plants. The numbers of independent plants (n) scored for each population are listed in Supplementary Table 5.

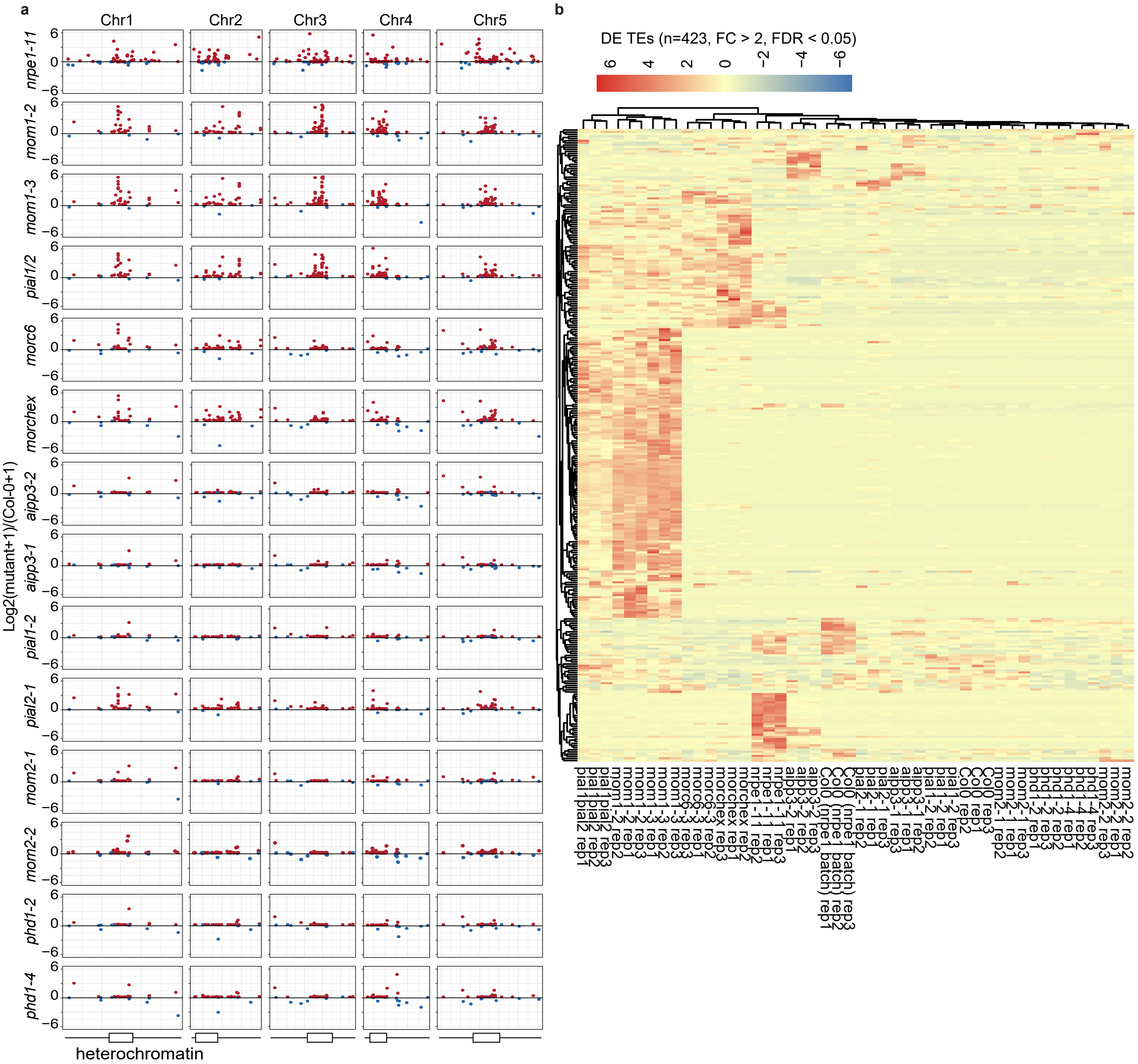


**Supplementary Fig. 6 | RNA-seq analysis of the mutants of MOM1 complex components. a,** Dotplots showing the differentially expressed TEs (compared to Col-0 control) over the five Arabidopsis chromosomes in the *nrpe1-11, mom1-2, mom1-3, pial1/2, morc6-3, morchex, aipp3-2, aipp3-1, pial1-2, pial2-1, mom2-1, mom2-2, phd1-2* and *phd1-4* mutant backgrounds. Red and blue dots indicate upregulated and down regulated TEs in mutants compared to Col-0 control, respectively. The positions of pericentromeric heterochromatin regions of each chromosome are annotated at the bottom of each plot. **b**, Heatmap showing the expression level of differentially expressed TEs (DE TEs, n=423) in three replicates of *mom1-2, mom1-3, pial1/2, morc6-3, morchex, aipp3-1, aipp3-2, pial1-2, pial2-1, mom2-1, mom2-2, phd1-2* and *phd1-4* mutant plants versus Col-0 plants. Expression level of these TEs in *nrpe1-11* mutant and corresponding Col-0 control plants are also plotted for comparison.
